# A connection between the ribosome and two *S. pombe* tRNA modification mutants subject to rapid tRNA decay

**DOI:** 10.1101/2023.09.18.558340

**Authors:** Thareendra De Zoysa, Alayna C. Hauke, Nivedita R. Iyer, Erin Marcus, Sarah M. Ostrowski, Justin C. Fay, Eric M. Phizicky

**Affiliations:** Department of Biochemistry and Biophysics, Center for RNA Biology, University of Rochester School of Medicine, Rochester, NY, USA 14642; Department of Biology, University of Rochester, Rochester, NY, USA 14627

**Keywords:** tRNA, 7-methylguanosine, 4-acetylcytidine, *S. pombe*, *S. cerevisiae*, TRM8, TAN1, rapid tRNA decay

## Abstract

tRNA modifications are crucial in all organisms to ensure tRNA folding and stability, and accurate translation in the ribosome. In both the yeast *Saccharomyces cerevisiae* and the evolutionarily distant yeast *Schizosaccharomyces pombe*, mutants lacking certain tRNA body modifications (outside the anticodon loop) are temperature sensitive due to rapid tRNA decay (RTD) of a subset of hypomodified tRNAs. Here we show that for each of two *S. pombe* mutants subject to RTD, mutations in ribosomal protein genes suppress the temperature sensitivity without altering tRNA levels. Prior work showed that *S*. *pombe trm8Δ* mutants, lacking 7-methylguanosine, were temperature sensitive due to RTD and that one class of suppressors had mutations in the general amino acid control (GAAC) pathway, which was activated concomitant with RTD, resulting in further tRNA loss. We now find that another class of *S. pombe trm8Δ* suppressors have mutations in *rpl* genes, encoding 60S subunit proteins, and that suppression occurs with minimal restoration of tRNA levels and reduced GAAC activation. Furthermore, *trm8Δ* suppression extends to other mutations in the large or small ribosomal subunit. We also find that *S. pombe tan1Δ* mutants, lacking 4-acetylcytidine, are temperature sensitive due to RTD, that one class of suppressors have *rpl* mutations, associated with minimal restoration of tRNA levels, and that suppression extends to other *rpl* and *rps* mutations. However, although *S. pombe tan1Δ* temperature sensitivity is associated with some GAAC activation, suppression by an *rpl* mutation does not significantly inhibit GAAC activation. These results suggest that ribosomal protein mutations suppress the temperature sensitivity of *S. pombe trm8*Δ and *tan1*Δ mutants due to reduced ribosome concentrations, leading to both a reduced requirement for tRNA, and reduced ribosome collisions and GAAC activation. Results with *S. cerevisiae trm8Δ trm4Δ* mutants are consistent with this model, and fuel speculation that similar results will apply across eukaryotes.

## Introduction

tRNAs contain numerous post-transcriptional modifications. These tRNA modifications are highly conserved evolutionarily within each domain of life, and in some cases throughout all of the domains (de Crecy-Lagard et al. 2012; Jackman and Alfonzo 2013). Modifications impart important functional or structural properties to the tRNAs, and consequently, lack of any of a number of tRNA modifications results in growth defects in the budding yeast *Saccharomyces cerevisiae* and in neurological, mitochondrial, or other disorders in humans (Suzuki 2021; Phizicky and Hopper 2023). Modifications within the anticodon loop frequently affect the efficiency or fidelity of translation (Urbonavicius et al. 2001; Kothe and Rodnina 2007; Nedialkova and Leidel 2015; Grosjean and Westhof 2016; Rozov et al. 2016; Ranjan and Rodnina 2017), consistent with their interactions in the A-site of the ribosome. By contrast, modifications in the main tRNA body often impair the folding and/or stability of the tRNA (Helm et al. 1999; Kadaba et al. 2004; Alexandrov et al. 2006; Chernyakov et al. 2008).

The biology of body modifications in eukaryotes is best understood in *S. cerevisiae*, in which lack of any of several body modifications targets a subset of the hypomodified tRNAs to either or both of two quality control decay pathways, particularly at higher temperatures (see (Phizicky and Hopper 2023)). Lack of each of several modifications, alone or in combination with other body modifications, targets mature tRNAs for 5’-3’ exonucleolytic degradation by the rapid tRNA decay (RTD) pathway, which is catalyzed by Xrn1 and Rat1 and inhibited by lack of Met22 (Chernyakov et al. 2008; Dewe et al. 2012), due to the accumulation of the metabolite adenosine 3’,5’ bis-phosphate (Dichtl et al. 1997; Yun et al. 2018). Thus, *trm8Δ trm4Δ* mutants lack 7-methylguanosine at G_46_ (m^7^G_46_) and 5-methylcytidine (m^5^C) and are temperature sensitive due to RTD of tRNA^Val(AAC)^ (Alexandrov et al. 2006; Chernyakov et al. 2008). Similarly, both *tan1Δ trm44Δ* mutants (lacking 4-acetylcytidine at C_12_ (ac^4^C_12_) and 2’-O-methyluridine at U_44_) and *trm1Δ trm4Δ* mutants (lacking N_2_,N_2_-dimethylguanosine at G_26_ (m^2,2^G_26_) and m^5^C) are temperature sensitive due to RTD of tRNA^Ser(CGA)^ and tRNA^Ser(UGA)^, and the temperature sensitivity of each of the *trm8Δ*, *trm1Δ*, and *tan1Δ* single mutants is associated with tRNA decay and inhibited by a *met22Δ* mutation (Chernyakov et al. 2008; Kotelawala et al. 2008; Dewe et al. 2012). By contrast, *trm6* mutants, lacking m^1^A58, are targeted by both the nuclear surveillance pathway and the RTD pathway. The nuclear surveillance pathway degrades pre-tRNAiMet lacking m^1^A58 using the 3’-5’ exonucleases Rrp6 and Rrp44 of the nuclear exosome, after oligoadenylation by Trf4 of the TRAMP complex (Kadaba et al. 2004; Kadaba et al. 2006; Wang et al. 2008), and in addition, the RTD pathway has a prominent role in decay of tRNAiMet lacking m^1^A58 in *trm6* mutants (Tasak and Phizicky 2022).

Our genetic analysis of the biology of body modification mutants in *S. pombe* has revealed strong evidence for conservation of the use of the RTD pathway in monitoring tRNA quality across at least the 600 million years separating *S. pombe* and *S. cerevisiae*, which pre-dates the Cambrian explosion during which the major animal phyla emerged (Parfrey et al. 2011). Thus, we showed that *S. pombe trm8Δ* mutants, lacking m^7^G46, were temperature sensitive due to reduced levels of tRNA^Tyr(GUA)^, and to some extent tRNA^Pro(AGG)^, and that each of four *trm8Δ* suppressors with restored tRNA levels had mutations in the *RAT1* ortholog *dhp1* (De Zoysa and Phizicky 2020). Similarly, we found that the temperature sensitivity of *S. pombe trm6Δ* mutants, lacking m^1^A58, was due to the decay of tRNAiMet by the RTD pathway, and not the nuclear surveillance pathway, based on isolation of multiple *trm6Δ* suppressors with mutations in *dhp1* or *tol1* (the *MET22* ortholog), and the lack of *trm6Δ* suppression by mutation of the *TRF4* ortholog *cid14* (Tasak and Phizicky 2022).

Our additional analysis of *S. pombe trm8Δ* suppressors also led to the discovery of a connection between the onset of RTD and activation of the general amino acid control (GAAC) pathway, equivalent to the integrated stress response pathway in humans. Thus, each of seven *S. pombe trm8Δ* suppressors that were sensitive to the GAAC activator 3-aminotriazole (3-AT) had null mutations in one of three components of the GAAC pathway *(gcn1^+^, gcn2^+^*, or *gcn3^+^* (*tif221^+^*)) and had partially restored levels of tRNA^Tyr^ and tRNA^Pro(AGG)^, consistent with the interpretation that decay of tRNA^Tyr^ and tRNA^Pro(AGG)^ in *S. pombe trm8Δ* mutants triggers GAAC activation, causing a further loss of these tRNAs. In support of this interpretation, we found that the lowest temperature at which tRNA decay was detected in *S. pombe trm8Δ* mutants coincides with the temperature at which a growth defect was first observed and GAAC activation was first evident (De Zoysa and Phizicky 2020). Moreover, the connection between the onset of RTD and GAAC activation extends to *S. cerevisiae*, as the lowest temperature at which the well-studied *S. cerevisiae trm8Δ trm4Δ* mutant showed a growth defect was accompanied by significant tRNA^Val(AAC)^ decay and GAAC activation. However, GAAC activation has opposite consequences in *S. cerevisiae trm8Δ trm4Δ* mutants, compared to *S. pombe trm8Δ* mutants, as introduction of GAAC mutations led to exacerbated temperature sensitivity and a further reduction in tRNA^Val(AAC)^ levels, consistent with the GAAC pathway acting to prevent further tRNA loss in *S. cerevisiae* (De Zoysa and Phizicky 2020).

Here, we show the existence of a connection between tRNA modification biology in *S. pombe* and proteins in the ribosome, which also appears to extend to *S. cerevisiae*. We find that a prominent class of spontaneous suppressors of the temperature sensitivity of *S. pombe trm8Δ* mutants have mutations in *rpl* genes, encoding proteins in the large ribosomal subunit, and that suppression is accompanied by little or no restoration of tRNA levels and reduced activation of the GAAC pathway. This suppression of *S. pombe trm8Δ* mutants extends to each of several tested *rpl* and *rps* genes (encoding proteins in the small subunit). Furthermore, the connection between tRNA modification and proteins in the ribosome also extends to another *S. pombe* body modification mutant. Thus, we find that *S. pombe tan1Δ* mutants, lacking ac^4^C12 in their tRNAs, are temperature sensitive due to RTD of two tRNAs, and that a large class of *tan1Δ* suppressors have mutations in ribosomal protein genes. However, *S. pombe tan1Δ* mutants have reduced GAAC activation, compared to *S. pombe trm8Δ* mutants, and an *rpl502Δ* mutation does not dramatically inhibit GAAC activation, but suppresses the *S. pombe tan1Δ* growth defect more efficiently than a *gcn2Δ* mutation. In contrast, we find that *rpl* and *rps* mutations exacerbate the temperature sensitivity of the well-studied *S. cerevisiae trm8Δ trm4Δ* mutant, known to be subject to RTD. We interpret these results in terms of a model in which *rpl* and *rps* mutations lead to reduced ribosome concentrations (Steffen et al. 2012; Cheng et al. 2019), resulting in reduced ribosome collisions and consequent GAAC activation (Yan and Zaher 2021; Kim and Zaher 2022), as well as in a reduced need for tRNA. Based on these results, we speculate that mutations in ribosomal protein genes will similarly act to influence growth defects of other modification mutants in *S. pombe* and *S. cerevisiae*, as well as in other eukaryotes.

## Results

### A class of *trm8***Δ** suppressors have mutations in *rpl* genes and distinct growth properties

To further understand mechanisms mediating the temperature sensitivity of *S. pombe trm8*Δ mutants, we identified additional suppressors with growth phenotypes different from those with mutations in the RTD pathway or the GAAC pathway, and then sequenced their genomes. We tested sensitivity to 5-fluorouracil (5-FU) for two reasons. First, our previous work showed that *S. pombe trm8Δ* mutants were modestly sensitive to 5-FU at higher temperatures (De Zoysa and Phizicky 2020), consistent with the 5-FU sensitivity of *S. cerevisiae trm8Δ* mutants (Gustavsson and Ronne 2008). Second, we had shown that *S. pombe trm8Δ dhp1* mutants were 5-FU resistant (De Zoysa and Phizicky 2020), likely because the *dhp1* mutations protect tRNAs from decay due to the lack of both m^7^G, and modifications inhibited by 5-FU (Frendewey et al. 1982; Santi and Hardy 1987; Huang et al. 1998). We also tested 3-AT sensitivity because *S. pombe trm8Δ gcn2* mutants, like other strains with mutations in the GAAC pathway, are sensitive to 3-AT, a competitive inhibitor of His3 that leads to histidine starvation and activation of the GAAC pathway, resulting in massive re-programming of gene expression that includes increased biosynthesis of His3 (Hinnebusch 2005).

In this way, we identified a group of four *trm8Δ* suppressors that suppressed *trm8Δ* temperature sensitivity in rich (YES) and complete minimal media lacking histidine (EMMC-His) almost as efficiently as a *trm8Δ* suppressor with a representative *dhp1* mutation (*trm8Δ dhp1-W326L*) or GAAC mutation (*trm8*Δ *gcn2-M1I*) (De Zoysa and Phizicky 2020). However, unlike the *dhp1* suppressors, these suppressors grew poorly on media containing 5-FU at 33°C, and unlike the GAAC suppressors, these suppressors were resistant to 3-AT at 37°C (Fig. 1A and Fig. S1). This class of *trm8Δ* suppressors all proved to have mutations in *rpl* genes (*rpl1701-Q72X, rpl502-Y44X, rpl1502-X201E,* and *rpl1102-K93X*), encoding different proteins (Rpl17, Rpl5, Rpl15, and Rpl11) in the 60S subunit of the ribosome (Table S1). Based on Pombase, all of these mutations are in one of two paralogous genes and are not essential, and the three mutant alleles with a premature stop codon are predicted to be null mutants.

**Fig. 1.**
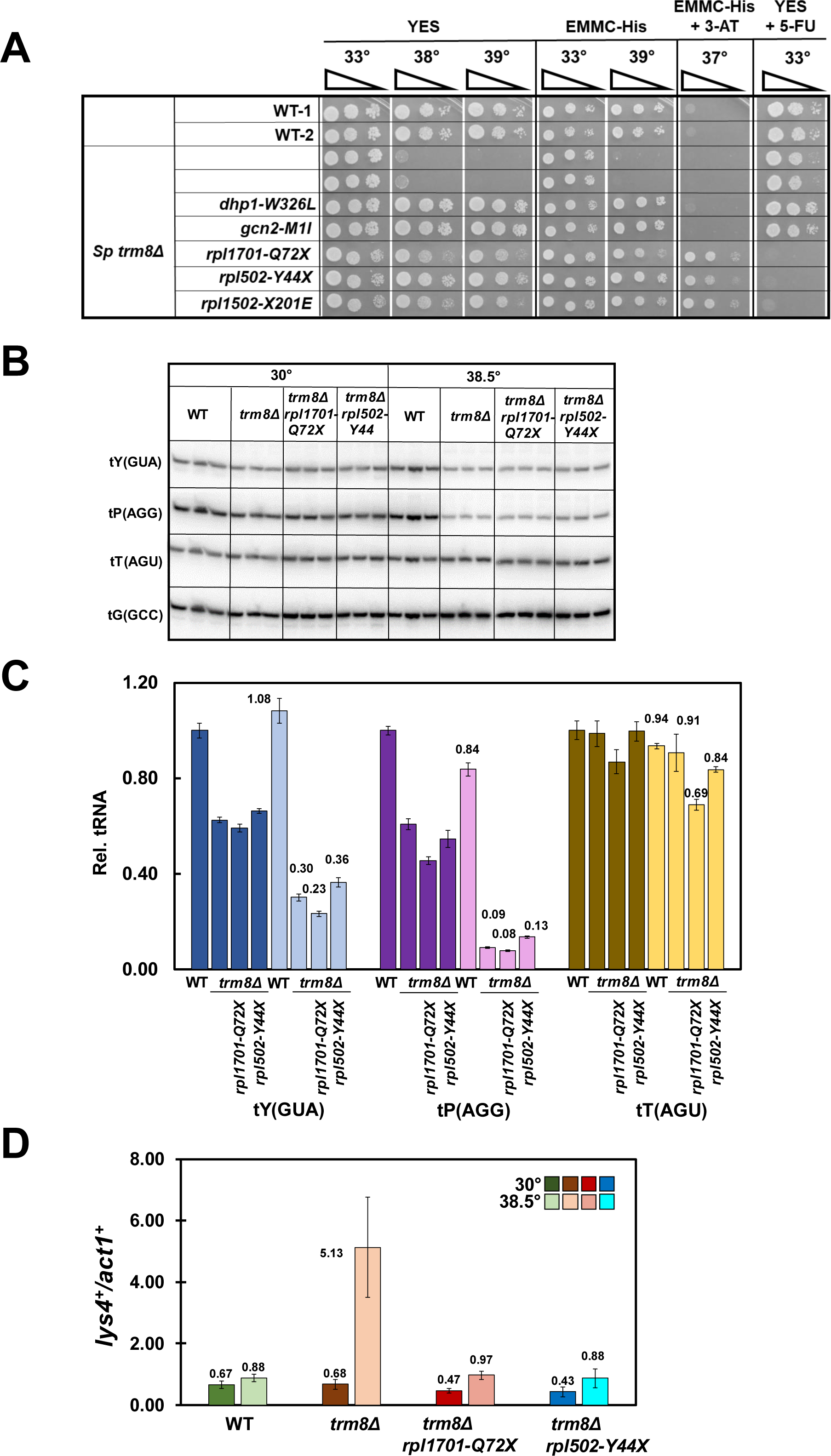
A class of *S. pombe trm8Δ* suppressors have mutations in *rpl* genes, minimally restore tRNA levels, and prevent activation of the GAAC pathway. **(A). *S. pombe trm8Δ* suppressors with distinctive growth properties have mutations in *rpl* genes.** *S. pombe trm8Δ* mutants and suppressors as indicated were grown overnight in YES media at 30°C, diluted to OD600 ∼ 0.5, serially diluted 10-fold in YES media, and 2 µl was spotted on plates containing YES, EMMC-His, EMMC-His with 10 mM 3-AT, and YES with 5-FU (30 µg/ml), as indicated. **(B). The *trm8Δ rpl1701-Q72X* and *trm8Δ rpl502-Y44X* suppressors do not appreciably prevent the decay of tRNA^Tyr(GUA)^ and tRNA^Pro(AGG)^ at 38.5 °C in EMMC-Leu media.** Strains transformed with a *leu2^+^* vector were grown in triplicate in EMMC-Leu media at 30°C, diluted into fresh media at 30°C and 38.5°C and grown for 10 hours, and then bulk RNA was analyzed for tRNA levels by northern blot analysis as described in Materials and Methods, with the indicated probes. **(C). Quantification of tRNA levels of WT, *trm8Δ*, and *trm8Δ rpl* mutants at 38.5°C in Fig. 1B**. The bar chart depicts relative levels of tRNA species at 30°C and 38.5°C, relative to their levels in the WT strain at 30 °C (each value itself first normalized to levels of the control non-Trm8 substrate tRNA^Gly(GCC)^). Standard deviations for each tRNA measurement are indicated. dark shades, 30°C growth; light shades, 38.5°C **(D). *S. pombe trm8Δ rpl1701-Q72X* and *trm8Δ rpl502-Y44X* mutants have reduced GAAC activation at 38.5°C, relative to *trm8Δ* strains.** Bulk RNA from the growth in Fig. 1B was used for RT-qPCR analysis of the mRNA levels of the GAAC target *lys4^+^* and the control *act1^+^*as described in Materials and Methods, and mRNA levels of *lys4^+^*were normalized to those of *act1^+^*.

To determine if the *rpl* mutations were responsible for the restoration of growth in the *trm8*Δ *rpl* suppressor mutants, we tested two of them for complementation by a [*leu2^+^*] plasmid containing the corresponding *rpl^+^* gene. Consistent with complementation, we found that, like the *trm8*Δ [*leu2^+^*] strain, expression of *rpl502^+^*in the *trm8*Δ *rpl502* mutant, or of *rpl1701^+^* in the *trm8*Δ *rpl1701* mutant (but not the vector controls) resulted in temperature sensitivity in EMMC-Leu media at 39°C, 5-FU resistance at 33°C, and 3-AT sensitivity at 38°C (Fig. S2).

### Suppression of the *S. pombe trm8*Δ temperature sensitivity by each of three *rpl* mutations occurs without significant restoration of tRNA levels

As the temperature sensitivity of *S. pombe trm8*Δ mutants is due to decay of tRNA^Tyr(GUA)^, and to a limited extent tRNA^Pro(AGG)^ (De Zoysa and Phizicky 2020), we analyzed tRNA levels of *trm8*Δ *rpl* suppressors after temperature shift from 30°C to 38.5°C. As expected, in *trm8*Δ mutants at 38.5°C, tRNA^Tyr(GUA)^ levels were reduced, to 30% of WT levels (Fig. 1B, 1C). Surprisingly, we found that in the *trm8*Δ *rpl1701-Q72X* suppressor the tRNA^Tyr(GUA)^ levels were not restored, but were if anything slightly reduced, relative to the *trm8*Δ mutant (23% vs 30%). Similarly, the tRNA^Pro(AGG)^ levels of the *trm8*Δ *rpl1701-Q72X* mutant (8%) were similar to, or slightly less, than those of the *trm8*Δ mutant (9%). Thus, for the *trm8*Δ *rpl1701-Q72X* suppressor, suppression is occurring without a detectable increase in levels of tRNA^Tyr(GUA)^ or of tRNA^Pro(AGG)^. Comparable analysis shows that the *trm8*Δ *rpl502-Y44X* suppressor had slightly increased levels of tRNA^Tyr(GUA)^ (36% vs 30%) and of tRNA^Pro(AGG)^ (13% vs 9%) relative to WT (Fig. 1B, 1C), and that the *trm8*Δ *rpl1102-K93X* suppressor also had little effect on levels of tRNA^Tyr(GUA)^ and tRNA^Pro(AGG)^ (Fig. S3). By contrast, our prior results showed that each of four independent *trm8Δ dhp1* suppressors invariably restored tRNA^Tyr(GUA)^ levels by more than 2.8-fold (2.8 to 3.9 fold) relative to those in the *trm8*Δ mutant, to more than 72% (73 - 85%) of WT levels, whereas GAAC mutations partially restored tRNA^Tyr(GUA)^ levels (by 1.7 to 2.1 fold), to 35 - 58% of WT levels (De Zoysa and Phizicky 2020). Thus, we conclude that as a group, these three *rpl* mutations suppress the temperature sensitivity of *S. pombe trm8*Δ mutants without obvious restoration of tRNA^Tyr(GUA)^ levels, or of tRNA^Pro(AGG)^ levels.

### Mutations in *rpl* genes prevent GAAC activation of *S. pombe trm8Δ* mutants

As we had previously shown that the temperature sensitivity of *S. pombe trm8*Δ mutants coincided with activation of the GAAC pathway and further loss of tRNA^Tyr(GUA)^ (De Zoysa and Phizicky 2020), we measured GAAC activation in two of the *S. pombe trm8*Δ *rpl* mutants (*trm8*Δ *rpl1701-Q72X* and *trm8*Δ *rpl502-Y44X*) at 38.5°C, using the same bulk RNA that we used for tRNA analysis in Fig. 2A and 2B. Consistent with our previous analysis (De Zoysa and Phizicky 2020), *S. pombe trm8*Δ mutants at 38.5°C had a 5.8-fold increase in relative mRNA levels of the GAAC target *lys4^+^* (normalized to the control *act1^+^*mRNA), compared to that from WT cells at 38.5°C (5.13 vs 0.88) and a 7.5-fold increase compared to that from *trm8*Δ mutants at 30°C (5.13 vs 0.68). By contrast, *trm8*Δ *rpl1701-Q72X* and *trm8*Δ *rpl502-Y44X* mutants had near baseline relative *lys4^+^*expression at both temperatures (0.97 and 0.88 vs 0.88 for WT, at 38.5°C) (Fig. 1D). Similar results were obtained by measuring relative *aro8^+^* mRNA levels, although the GAAC activation is not as strong (Fig. S4), as we observed previously (De Zoysa and Phizicky 2020). We conclude that the *rpl* mutations inhibit the GAAC activation normally observed in *trm8*Δ mutants at 38.5°C, likely due to reduced ribosome concentrations arising from mutation of one of the two paralogs of each ribosomal protein.

**Fig. 2.**
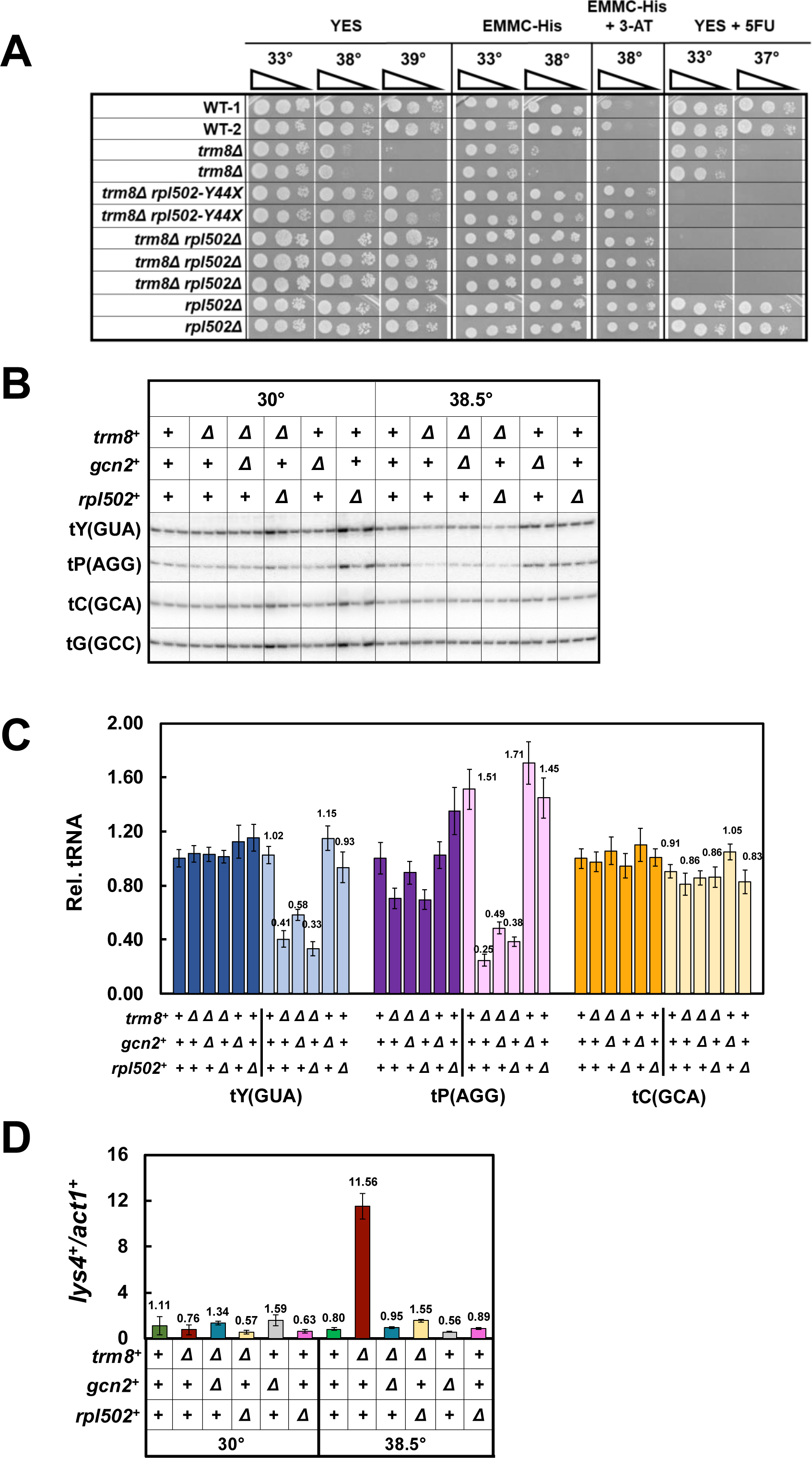
A reconstructed *trm8Δ rpl502Δ* mutant has the same properties as the *trm8Δ rpl502-Y44X* mutant. **(A). A reconstructed *trm8Δ rpl502Δ* mutant suppresses the *S. pombe trm8Δ* growth defect as efficiently as the original *trm8Δ rpl502-Y44X* mutant.** WT, *trm8Δ*, *trm8Δ rpl502-Y44X*, and each of three independent reconstructed *trm8Δ rpl502Δ* strains and two independent *rpl502Δ* strains were analyzed for growth on media containing YES, EMMC-His, EMMC-His with 10 mM 3-AT, and YES with 5-FU (30 µg/ml), as indicated. **(B). A reconstructed *trm8Δ rpl502Δ* suppressor does not substantially prevent decay of tRNA^Tyr(GUA)^ and tRNA^Pro(AGG)^ in *trm8Δ* mutants at 38.5°C**. Strains were grown in YES media at 30°C, shifted to 38.5°C for 9 hours, and analyzed for tRNA levels as described in Fig. 1B. **(C). Quantification of tRNA levels of WT, *trm8Δ* and *trm8Δ rpl502Δ* mutants at 38.5°C. (D). A reconstructed *trm8Δ rpl502Δ* suppressor inhibits GAAC activation observed in *trm8Δ* mutants in YES Media at 38.5°C**. Bulk RNA from the growth in Fig. 2B was analyzed for GAAC activation as described in Fig. 1D.

A reconstructed *S. pombe trm8Δ rpl502Δ* mutant has the same properties as a *trm8*Δ *rpl502-Y44X* mutant.

To ensure that the effects of the *trm8*Δ *rpl502-Y44X* mutant were solely due to the *rpl502* mutation, rather than some background mutation that arose in the original *trm8Δ* strain before selection for suppressors, we introduced an *rpl502Δ* mutation into a clean *trm8Δ* mutant and tested the reconstructed *trm8Δ rpl502Δ* mutant. We found that a reconstructed *trm8*Δ *rpl502*Δ strain suppressed the growth defect of a *trm8Δ* mutant almost identically to the *trm8*Δ *rpl502-Y44X* mutant, with similar growth in YES and EMMC media at high temperature, and similar 5-FU sensitivity and 3-AT resistance (Fig. 2A). Also like *trm8*Δ *rpl502-Y44X* mutants, *trm8*Δ *rpl502*Δ mutants did not restore tRNA^Tyr(GUA)^ levels after temperature shift to 38.5°C in YES media (33% vs 41% for *trm8*Δ mutants) whereas levels of tRNA^Tyr(GUA)^ were modestly restored in *trm8Δ gcn2Δ* mutants (58% vs 41%), as we observed previously (De Zoysa and Phizicky 2020) (Fig. 2B, 2C). Similarly, tRNA^Pro(AGG)^ levels were not efficiently restored in *trm8*Δ *rpl502*Δ mutants (38%) relative to 25% in *trm8*Δ mutants and 49% in *trm8Δ gcn2Δ* strains. Furthermore, like *trm8*Δ *rpl502-Y44X* mutants, *trm8*Δ *rpl502*Δ mutants efficiently suppressed the GAAC activation observed in *trm8Δ* mutants, measured by relative *lys4^+^* levels (from a 14.5-fold increase in *trm8Δ* mutants, to a 1.9-fold increase in *trm8*Δ *rpl502*Δ mutants), compared to near baseline 1.2-fold increase in a *trm8Δ gcn2Δ* strain (Fig. 2D; Fig. S5). We conclude that the suppression effects of the *trm8*Δ *rpl502-Y44X* mutant are explicitly due to the *rpl* mutation, and infer that the suppression by the *trm8*Δ *rpl1701-Q72X* and *trm8Δ rpl1102-K93X* mutations is also due to the *rpl* mutations.

We note that, as the control *rpl502*Δ mutants are as resistant to 3-AT as the *trm8*Δ *rpl502Δ* mutants (Fig. 2A), this property is due to the *rplΔ* mutation and not the *trm8Δ* mutation in these strains.

### Deletion of any of several *rpl^+^* or *rps^+^* genes suppress the temperature sensitivity of *S. pombe trm8*Δ mutants

To determine if the spontaneous *S. pombe trm8Δ rpl* suppressors that we had found by selection represented a specific subset of ribosomal protein genes, we investigated whether or not suppression would extend to deletion of other *rpl^+^*or *rps^+^* genes. We focused on six ribosomal protein genes that had two copies and thus were not essential, including *rpl1701*, for which we already had a spontaneous *trm8Δ* suppressor. Deletion of each of the two *rps* genes (*rps2801Δ* and *rps802Δ*, encoding Rps28 and Rps8) and four *rpl* genes examined (*rpl1601Δ*, *rpl1202Δ, rpl2802Δ*, and *rpl1701Δ*, encoding Rpl16, Rpl12, Rpl28, and Rpl17) resulted in efficient suppression of the *trm8Δ* temperature sensitivity in EMMC-His media, although suppression was somewhat weaker for the *rpl1601Δ* mutation (Fig. 3A, 3B). Similarly, all five of the *trm8Δ rplΔ* and *trm8Δ rpsΔ* mutants that were robust *trm8Δ* suppressors were also sensitive on YES+5-FU media at 33°C, whereas the weak *trm8Δ rpl1601Δ* suppressor was slightly sensitive on YES+5-FU. We conclude that *rplΔ* and *rpsΔ* mutations generally cause suppression of the *S. pombe trm8Δ* growth defect, presumably as a consequence of the reduced concentrations of the corresponding ribosomal subunits (Warner 1999; Cheng et al. 2019).

**Fig. 3.**
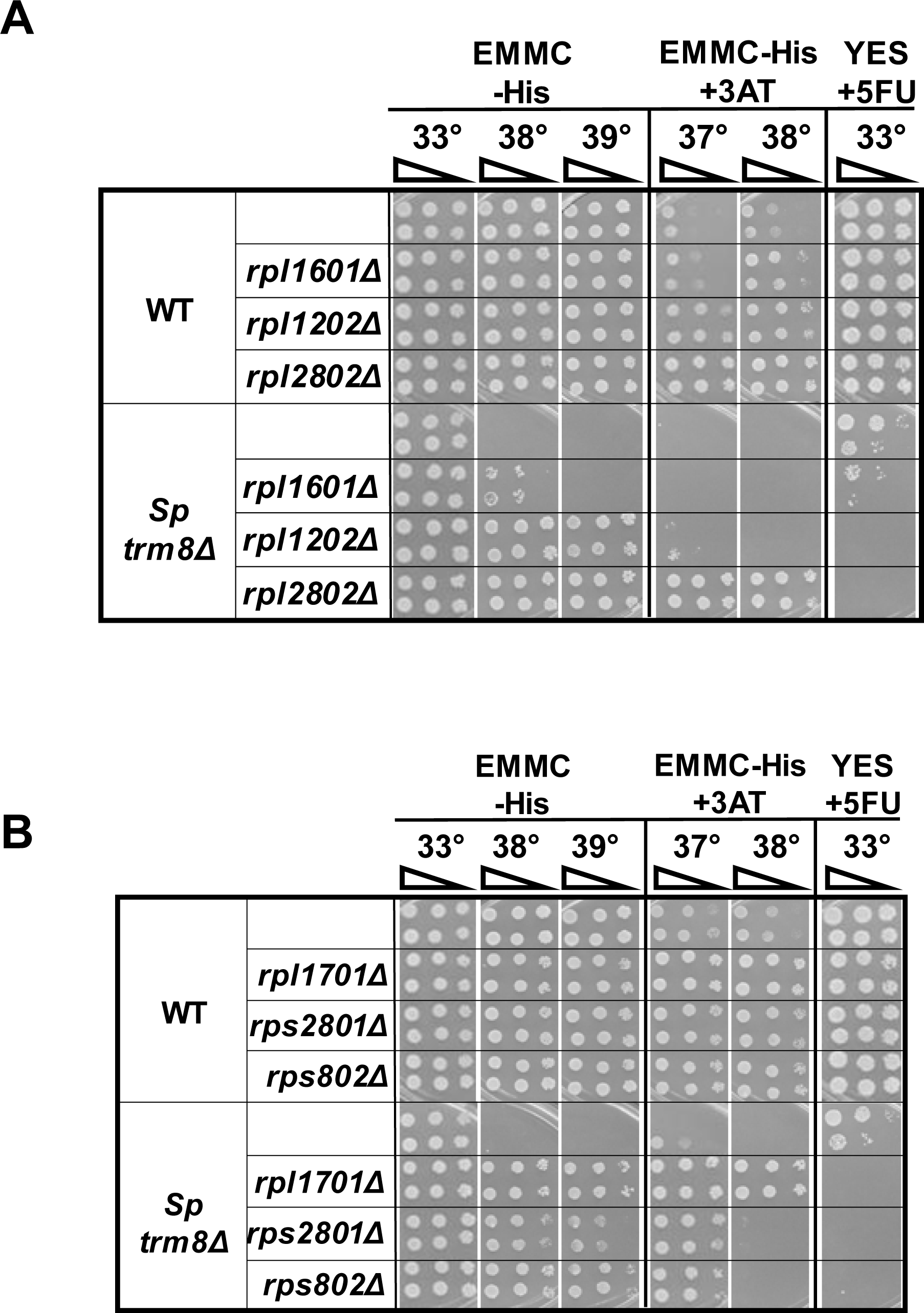
Deletion of any of several *rpl^+^* or *rps^+^* genes restores growth of *S. pombe trm8Δ* mutants at elevated temperatures. **(A, B).** *S. pombe rpl* genes and *rps* genes as indicated were deleted in WT and trm8Δ strains, and cultures were grown and analyzed on the indicated media.

By contrast, we found that 3-AT resistance was not correlated with the efficiency of suppression in *trm8Δ rplΔ* strain, as one of the robust *trm8Δ* suppressors was almost completely 3-AT sensitive (*trm8Δ rpl1202Δ*) and two other robust suppressors were only moderately 3-AT resistant (*trm8Δ rps2801Δ* and *trm8Δ rps802Δ*), whereas the robust *trm8Δ rplΔ2802Δ* and *trm8Δ rpl1701Δ* suppressor strains were strongly 3-AT resistant. This result is consistent with our finding above that 3-AT resistance is a property of the mutation in the ribosomal protein gene, and not the *trm8Δ* mutation, However, it is not immediately clear why some mutations in ribosomal protein genes result in 3-AT resistance and others result in sensitivity, although it is known that different mutations in ribosomal protein genes in *S. cerevisiae* result in different growth properties and phenotypes (Steffen et al. 2012; Cheng et al. 2019).

### An *S. pombe tan1Δ* mutant lacks 4-acetylcytidine and is temperature sensitive due to decay of tRNA^Leu(AAG)^ and tRNA^Leu(UAG)^ by the RTD pathway

To explore the generality of the effect of the ribosome and GAAC mutations on the growth of *S. pombe* modification mutants, we examined an *S. pombe tan1Δ* mutant. It is well established from work in *S. cerevisiae* that *tan1Δ* mutants lack 4-acetylcytidine at C12 (ac^4^C12) in their tRNA^Leu^ and tRNA^Ser^ species, which have a long variable arm (Johansson and Bystrom 2004), that *tan1Δ trm44Δ* mutants are temperature sensitive due to RTD of tRNA^Ser(CGA)^ and tRNA^Ser(UGA)^ (Chernyakov et al. 2008), and that the temperature sensitivity of *tan1Δ* single mutants is associated with decay of tRNA^Ser(CGA)^ and tRNA^Ser(UGA)^ and is suppressed by a *met22Δ* mutation (Dewe et al. 2012). Indeed, we find that the temperature sensitivity of *S. cerevisiae tan1Δ* mutants is completely suppressed by overexpression of tRNA^Ser(CGA)^ and nearly as efficiently suppressed by overexpression of tRNA^Ser(UGA)^ (Fig. S6).

Consistent with the biology of Tan1 in *S. cerevisiae*, we find that *S. pombe tan1Δ* mutants are temperature sensitive due to decay of two tRNA species by the RTD pathway. As anticipated based on the conservation of Tan1 sequence and the Tan1 requirement for ac^4^C12 modification of tRNA^Leu^ and tRNA^Ser^ within a CCG motif (Aravind and Koonin 2001; Sharma et al. 2015; Sas-Chen et al. 2020; Broly et al. 2022), purified tRNA^Leu(CAA)^ from *S. pombe tan1Δ* mutants lacks any detectable ac^4^C, compared to that in WT cells, but has similar levels of four control modifications (Fig. 4A). In addition, we find that *S. pombe tan1Δ* mutants are temperature sensitive on both YES media and EMMC-Leu media at 38°C (Fig. 4B). The *S. pombe tan1Δ* temperature sensitivity is associated with reduced levels of tRNA^Leu(AAG)^ and tRNA^Leu(UAG)^ at 30°C that are further reduced at 39°C (relative to those in WT cells), whereas levels of each of the other tRNA^Leu^ and tRNA^Ser^ species do not conform to this pattern (Fig. 4C, Fig. S7, S8). Furthermore, the temperature sensitivity of *S. pombe tan1Δ* mutants is almost entirely due to tRNA^Leu(AAG)^, and to some extent tRNA^Leu(UAG)^, as overexpression of these two tRNAs, or tRNA^Leu(AAG)^ alone, efficiently suppresses the temperature sensitivity (Fig. 4D). Moreover, as anticipated from Tan1 biology in *S. cerevisiae*, whole genome sequencing resulted in identification of three *S. pombe tan1Δ* suppressors with *dhp1* mutations (*dhp1-I196T*, *dhp1-P677S*, and *dhp1-P203L*), indicating decay of these tRNAs by the RTD pathway; these suppressors will be discussed elsewhere.

**Fig. 4.**
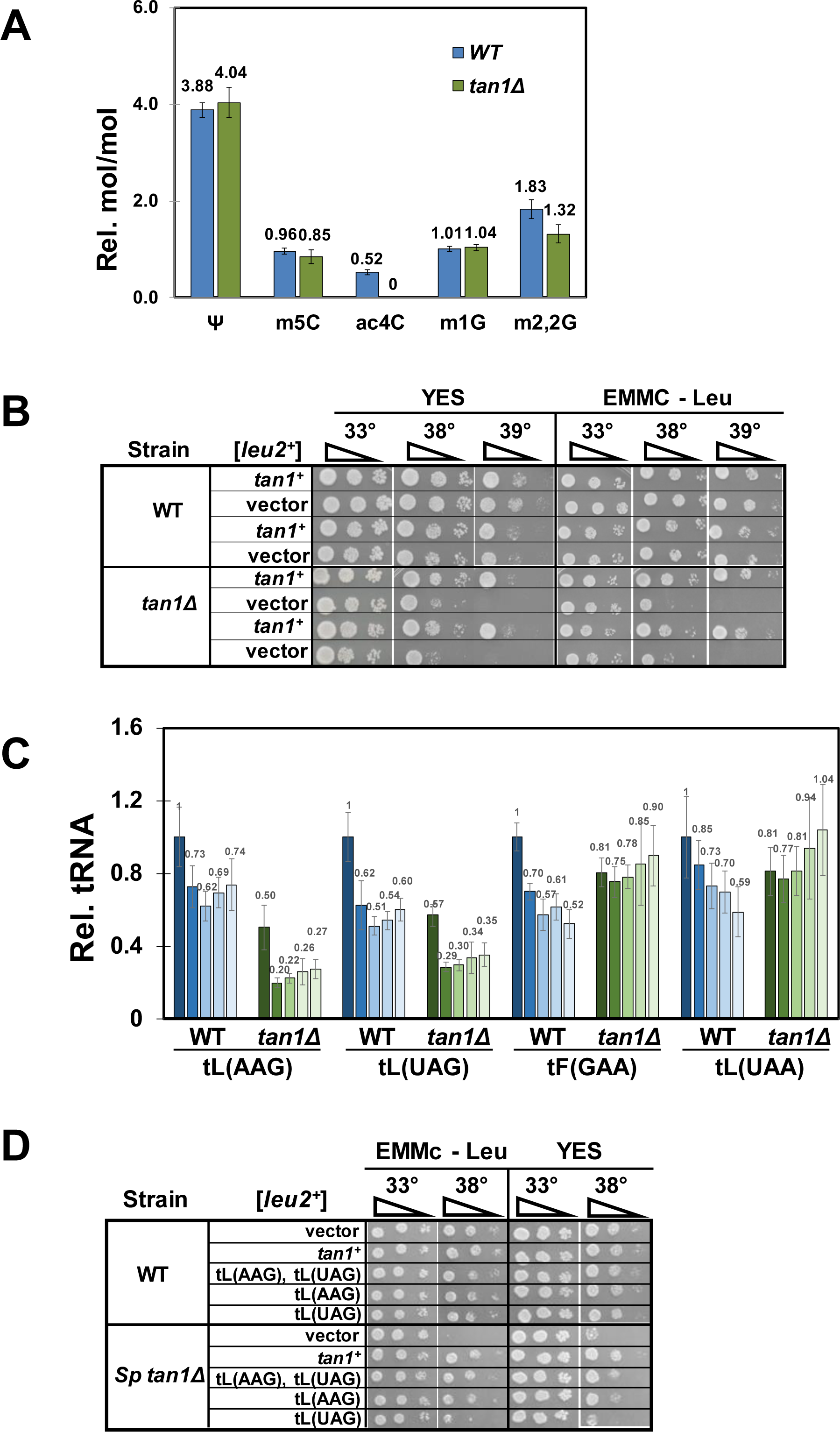
*S. pombe tan1Δ* mutants lack 4-acetylcytidine and are temperature sensitive due to decay of tRNA^Leu(AAG)^ and tRNA^Leu(UAG)^. **(A).** *S. pombe tan1Δ* mutants lack ac^4^C in their tRNA^Leu(CAA)^. *S. pombe* WT and *tan1Δ* mutants were grown in triplicate in YES media at 30°C to late log phase, and tRNA^Leu(CAA)^ was purified from bulk RNA, digested to nucleosides, and analyzed for modifications by HPLC as described in Materials and Methods. **(B). *S. pombe tan1Δ* mutants are temperature sensitive.** *S. pombe* WT and *tan1Δ* mutants were transformed with a [*leu2^+^*] or a [*leu2^+^ tan1^+^*] plasmid as indicated, and transformants were grown overnight in EMMC-Leu media, serially diluted, and plated on media as indicated. **(C). *S. pombe tan1Δ* mutants have reduced levels of tRNA^Leu(AAG)^ and tRNA^Leu(UAG)^ at 39°C.** *S. pombe* WT and *tan1Δ* mutants were transformed with a [*leu2^+^*] plasmid, and transformants were grown in EMMC-Leu media to mid-log phase, diluted and shifted to 39°C and grown for 12 hours, and RNA was isolated at 0, 3, 6, 9, and 12 hours and analyzed by Northern blot as described in Materials and Methods, with the indicated probes. shades of blue, WT strains, analyzed at 0 (darkest) through 12 hours (lightest); shades of green, *tan1Δ* strains **(D). The temperature sensitivity of *S. pombe tan1Δ* mutants is efficiently suppressed by overexpression of tRNA^Leu(AAG)^ and, to some extent, tRNA^Leu(UAG)^.** *S. pombe* WT and *tan1Δ* mutants were transformed with a [*leu2^+^*] plasmid expressing tRNAs as indicated, or a vector control, and transformants were grown overnight in EMMC - Leu media at 30 °C and analyzed for growth, as in Fig 1A, on plates containing EMMC – Leu or YES media.

### A prominent class of *S. pombe tan1Δ* suppressors have mutations in *rpl* genes or related genes predicted to reduce 60S subunits

Consistent with our results with *S. pombe trm8Δ* suppressors, we find that several spontaneous suppressors of the temperature sensitivity of *S. pombe tan1Δ* mutants have mutations in ribosomal protein genes. Thus, of seven *tan1Δ* suppressors that are resistant to 3-AT at 35°C and somewhat sensitive to 5-FU at 38°C, whole genome sequencing showed that five have *rpl* mutations (*rpl1101-N7fs*, *rpl1102-R133fs*, *rpl1502-R67_R68_insVR*, *rpl1701-T35fs*, *rpl2802-G113fs*, and *rpl3001-N80fs*) and the seventh has a *grn1-N37fs* mutations. *S. pombe* Grn1 and its *S. cerevisiae* ortholog Nug1 are critical GTPases required for pre-60S rRNA processing (Bassler et al. 2001; Du et al. 2006). It is therefore clear that this selection (temperature resistance) and screening procedure (3-AT-resistance, 5-FU sensitivity) is consistently yielding *tan1Δ* suppressors with mutations predicted to reduce the amount of the 60S subunit, just as we observed with *trm8Δ* suppressors.

We confirmed two of the ribosomal protein mutations as *tan1Δ* suppressors by introduction of the corresponding deletion mutation into a clean *tan1Δ* background. Thus, we find that an *rpl2802Δ* mutation efficiently suppresses the growth phenotypes of an *S. pombe tan1Δ* mutant, resulting in temperature resistance on EMMC-His media and YES media at 38°C, 3-AT resistance at 35°C, and 5-FU sensitivity at 35°C (Fig. 5A). Similarly, an *rpl1701Δ* mutation also efficiently suppresses each of the growth phenotypes of an *S. pombe tan1Δ* mutant, resulting in temperature resistance, 3-AT resistance, and 5-FU sensitivity (Fig. 5B).

**Fig. 5.**
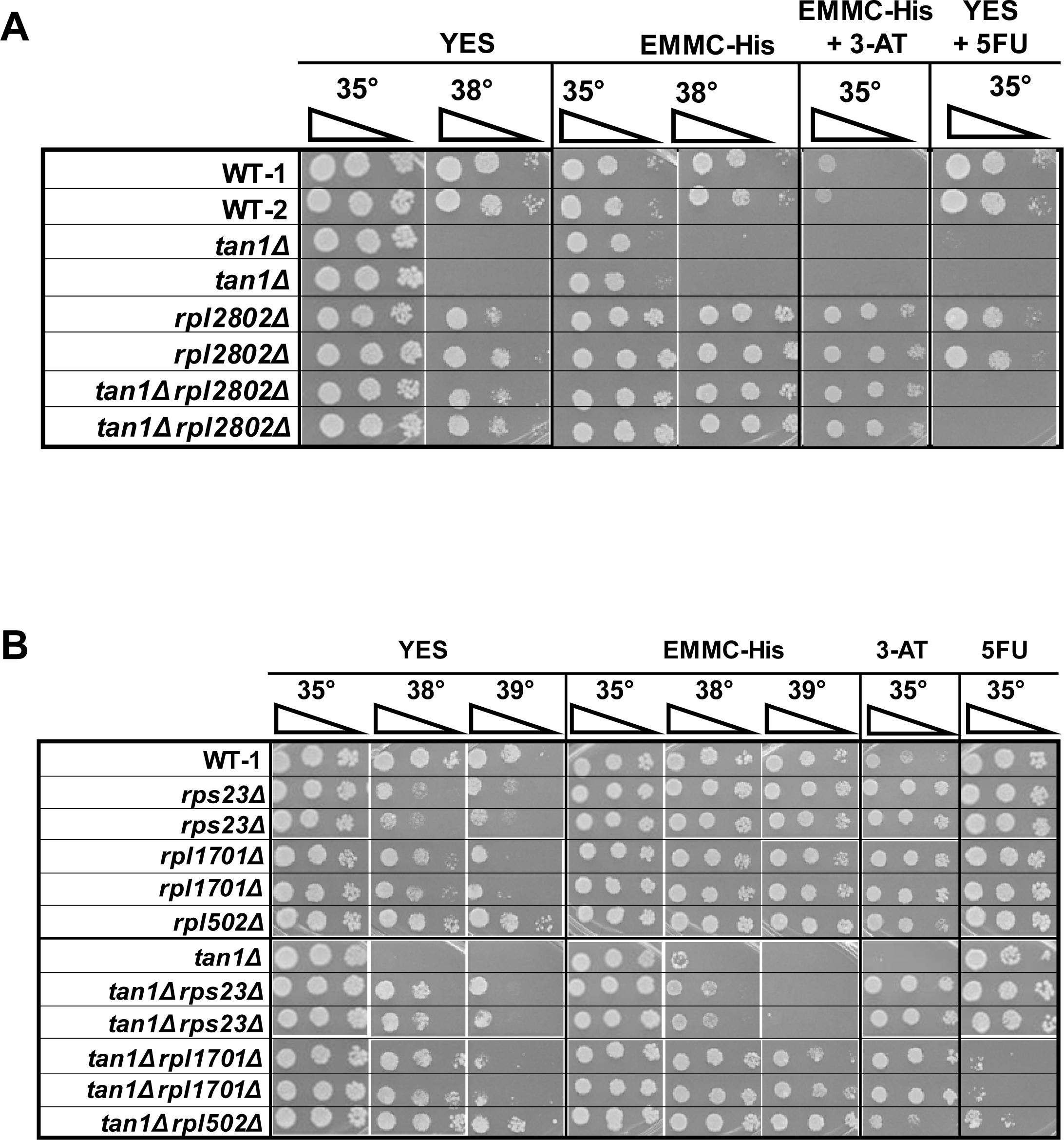
The temperature sensitivity of *S. pombe tan1Δ* mutants is efficiently suppressed by *rplΔ* and *rpsΔ* mutations. **(A). A reconstructed *tan1Δ rpl2802Δ* mutant suppresses the *S. pombe tan1Δ* growth defect.** The *rpl2802Δ* mutation was introduced into WT, and *tan1Δ* strains, and then WT, *tan1Δ*, *rpl2802Δ*, *tan1Δ rpl2802Δ* strains were grown overnight in YES media and analyzed for growth on media containing YES, EMMC - His, EMMC - His with 10 mM 3-AT, and YES with 5-FU (30 µg/ml), as indicated. **(B). A reconstructed *tan1Δ rpl1701Δ* mutant suppresses the *S. pombe tan1Δ* growth defect, as does each of two other tested deletions of ribosomal proteins.** The *rpl1701Δ*, *rps23Δ*, and *rpl502Δ* mutations were introduced into WT and *tan1Δ* strains, and then strains were grown overnight in YES media and analyzed for growth on media containing YES, EMMC - His, EMMC - His with 10 mM 3-AT, and YES with 5-FU (30 µg/ml), as indicated.

In addition, we find that *tan1Δ* suppression is not restricted to the specific ribosomal protein mutations that were selected, based on analysis of two other ribosomal protein genes. Thus, we find that an *rps23Δ* mutation and an *rpl502Δ* mutation each efficiently suppress the temperature sensitivity of the *tan1Δ* mutant, although we note that the *tan1Δ rps23Δ* mutant is not 5-FU sensitive, and that the *tan1Δ* r*pl502Δ* mutant is only slightly 3-AT resistant at 35°C (Fig. 5B).

### The *rpl502Δ* mutation suppresses the temperature sensitivity of the *S. pombe tan1Δ* mutant without restoring tRNA levels or affecting GAAC activation

To assess the role of the *rpl502Δ* mutation in suppression of the *S. pombe tan1Δ* growth defect, we grew cells at 30°C in YES media, shifted to 39°C, and analyzed both tRNA levels and GAAC activation after 9 hours. As we saw for the *trm8Δ rpl502Δ* strain, we find that the *tan1Δ rpl502Δ* strain does not have restored tRNA levels after growth at 39°C (Fig. 6A, 6B). Relative tRNA^Leu(AAG)^ levels are reduced substantially in *tan1Δ* strains after growth at 39°C (to 24% of WT), and remain low in the *tan1Δ rpl502Δ* strain at 39°C (19% of WT), but each of two control tRNAs are unaffected in both strains. However, we find that although an *S. pombe tan1Δ* mutant activates the GAAC pathway, based on analysis of relative *lys4^+^*mRNA levels, the activation is weak, and is only modestly reduced in a *tan1Δ rpl502Δ* strain (Fig. 6C). Thus, we find that a *tan1Δ* mutant has a 3.3-fold increase in relative *lys4^+^* mRNA levels at 39°C (1.62) compared to WT at 39°C (0.49) and a 2.1-fold increase compared to *tan1Δ* mutants at 30°C (0.80). This is substantially weaker than the GAAC activation of a *trm8Δ* mutant in the same experiment, which increases 23.4-fold compared to WT at 39°C (11.45 vs 0.49) and 12.6-fold, compared to *trm8Δ* mutants at 30°C (11.45 vs 0.91). Moreover, we find that an *rpl502Δ* mutation has only a minimal effect on GAAC activation by a *tan1Δ* mutant. Thus, a *tan1Δ rpl502Δ* mutant has a 2.5-fold increase in relative *lys4^+^* levels at 39°C relative to 30°C (1.13 vs 0.46), compared to 2.1-fold for the *tan1Δ* mutant. and a 2.31-fold increase at 39°C relative to WT at 39°C (1.13 vs 0.49), compared to 3.31-fold for the *tan1Δ* mutant.

**Fig. 6.**
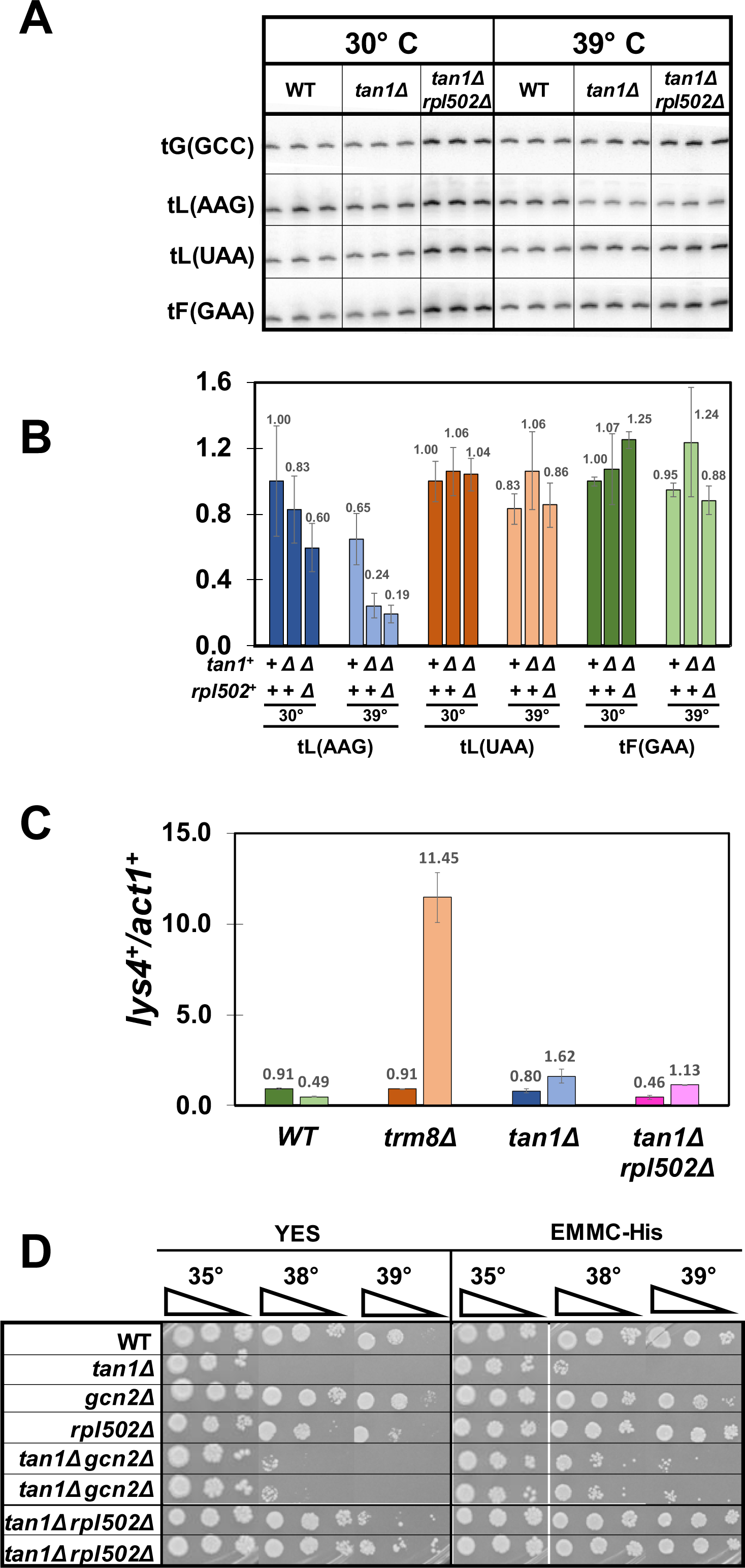
An *S. pombe tan1Δ rpl502Δ* suppressor does not rescue the tRNA^Leu(AAG)^ decay observed in *tan1Δ* mutants and only minimally inhibits the modest GAAC activation. **(A). An *S. pombe tan1Δ rpl502Δ* suppressor does not rescue the tRNA^Leu(AAG)^ decay observed in *tan1Δ* mutants.** Strains were grown in YES media at 30°C, shifted to 39°C for 9 hours, and bulk RNA was analyzed for tRNA levels as described in Fig. 1B. **(B). Quantification of tRNA levels of WT, *tan1Δ*, and *tan1Δ rpl502Δ* mutants at 39°C in Fig. 6A. (C). Analysis of GAAC activation.** Bulk RNA from the growth in Fig. 6A was analyzed for GAAC activation as described in Fig. 1D. **(D). The temperature sensitivity of a *tan1Δ* mutant is more efficiently suppressed in *tan1Δ rpl502Δ* strains than in *tan1Δ gcn2Δ* strains.** WT and *tan1Δ* strains with *rpl502Δ* or *gcn2Δ* mutations were grown overnight in YES media and analyzed for growth on media containing YES or EMMC - His, as indicated.

A similar analysis after growth of strains in EMMC-Leu media yields similar results regarding the effect of the *rpl502Δ* mutation on *S. pombe tan1Δ* mutants (Fig. S9). We observe reduced levels of tRNA^Leu(AAG)^ in the *S. pombe tan1Δ* mutant at 39°C (23% of WT levels at 30°C), which remain at similar or slightly lower levels in the *tan1Δ rpl502Δ* strain (19%). Although we observe some activation of the GAAC pathway in an *S. pombe tan1Δ* mutant, based on analysis of relative *lys4^+^* mRNA levels, the *rpl502Δ* mutation has only a modest effect on this activation. The *tan1Δ* mutant has a 3.8-fold increase in relative *lys4^+^* mRNA levels, compared to WT at 39°C (4.79 vs 1.27) (which is smaller than the 5.7-fold in the *trm8Δ* mutant), whereas the *tan1Δ rpl502Δ* strain has a 2.6 fold increase (Fig. S9).

These weak effects of the *tan1Δ* mutant and the t*an1Δ rpl502Δ* mutant on GAAC activation are consistent with the observation that an *rpl502Δ* mutation is a more efficient suppressor of the temperature sensitivity of an *S. pombe tan1Δ* mutant than is a *gcn2Δ* mutation, on either YES media or EMMC-His media at 38°C (Fig. 6D).

We infer from these results that the *rpl502Δ* mutation suppresses that *tan1Δ* mutant by reducing the need for tRNA^Leu(AAG)^, and likely tRNA^Leu(UAG)^, due to the reduced number of translating ribosomes arising from the *rpl502Δ* mutation, rather than by effects on the GAAC pathway due to reduced ribosome collisions.

### In *S. cerevisiae,* deletion of *rpl* or *rps* genes exacerbates the temperature-sensitivity of *trm8*Δ *trm4*Δ mutants

To investigate the evolutionary conservation of the effect of loss of ribosomal protein genes on mutants lacking m^7^G, we examined the growth properties arising from deletion of *RPL* or *RPS* genes in an *S. cerevisiae trm8*Δ *trm4*Δ mutant. This mutant is known to be highly temperature sensitive due to decay of tRNA^Val(AAC)^ by the RTD pathway (Alexandrov et al. 2006; Chernyakov et al. 2008), and like the *S. pombe trm8Δ* mutant, undergoes GAAC activation coincident with temperature sensitivity and the onset of tRNA decay (De Zoysa and Phizicky 2020). However, unlike the case with *S. pombe trm8Δ* mutants, *gcn2Δ* and *gcn1Δ* mutations each exacerbate the temperature sensitivity of *S. cerevisiae trm8Δ trm4Δ* mutants, and exacerbate the loss of tRNA^Val(AAC)^ at 32°C, leading to the conclusion that GAAC activation in *S. cerevisiae trm8Δ trm4Δ* mutants prevents further loss of tRNA^Val(AAC)^ and is beneficial (De Zoysa and Phizicky 2020).

We find that deletion of each of four tested *RPL* genes (*RPL1A, RPL11B, RPL17B,* or *RPL33B*) or three *RPS* genes (*RPS0A, RPS22A, RPS28A*) in the *S. cerevisiae trm8*Δ *trm4*Δ mutant does not suppress the temperature sensitivity of the strain, but rather, exacerbates its temperature sensitivity on rich (YPD) media and/or minimal (SD) media (Fig. 7A, 7B). We note that the exacerbated growth defect in the *S. cerevisiae trm8*Δ *trm4*Δ *rpl1AΔ* mutant is less distinct, which is likely related to the fact that although the tested ribosomal protein genes are all duplicated, the *RPL1A* gene is known to be the minor paralog (Petitjean et al. 1995; Steffen et al. 2008). Overall, the exacerbated growth defect in *S. cerevisiae trm8*Δ *trm4*Δ mutants bearing *rplΔ* or *rplsΔ* mutations is consistent with the opposite effects of GAAC activation in *S. pombe trm8Δ* mutants and *S. cerevisiae trm8*Δ *trm4*Δ mutants.

**Fig. 7.**
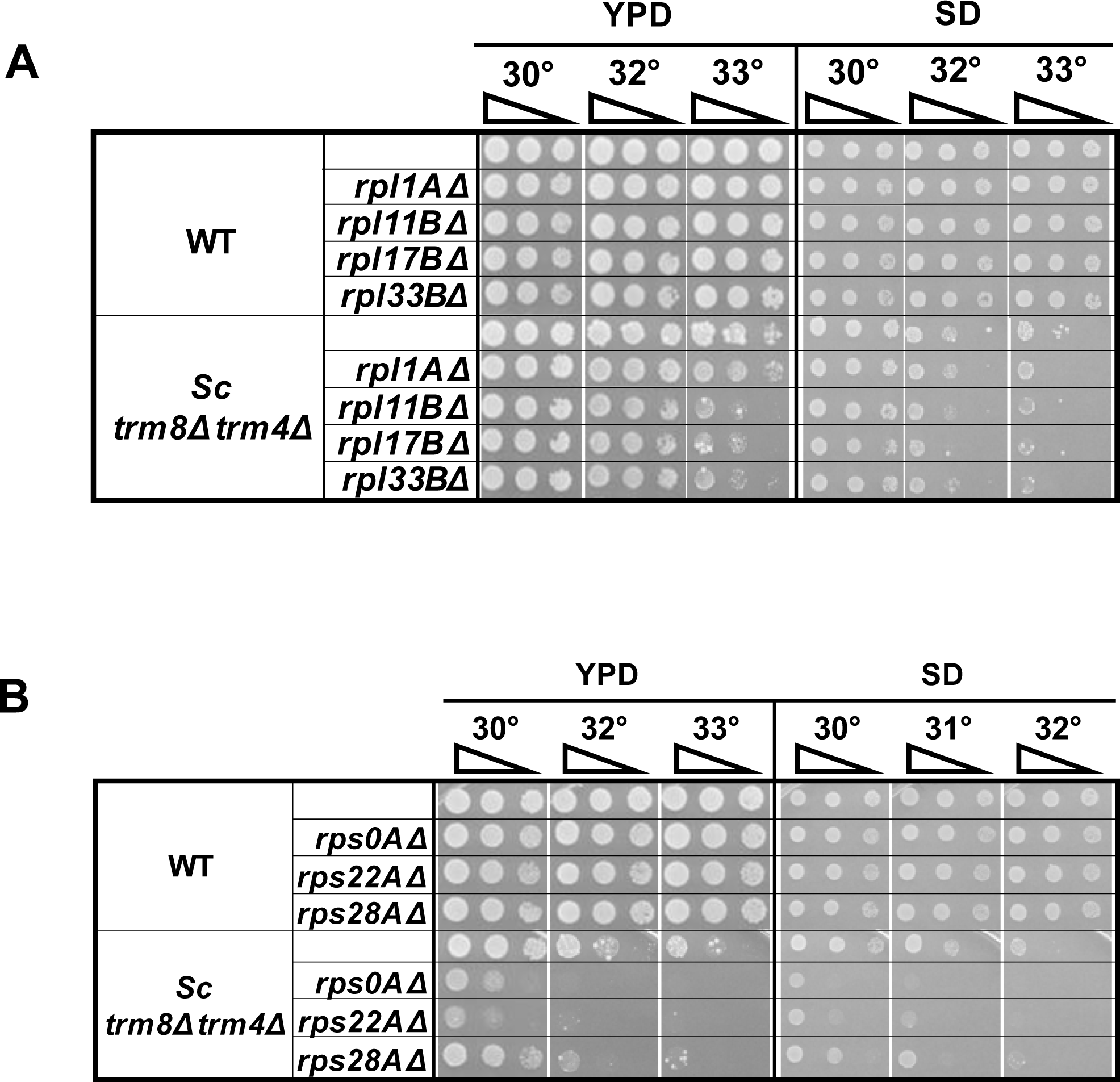
Deletion of any of several ribosomal protein genes exacerbates the temperature sensitivity of *S. cerevisiae trm8Δ trm4Δ* mutants in YPD Media. *rpl* genes **(A)** and *rps* genes **(B)** as indicated were deleted in *S. cerevisiae* WT and *trm8Δ trm4Δ* strains, and cultures were grown and analyzed on the indicated media.

## Discussion

We have provided evidence here that for each of two *S. pombe* body modification mutants (*trm8Δ* mutants and *tan1Δ* mutants), that are temperature sensitive due to RTD of specific hypomodified tRNAs, mutations in ribosomal protein genes suppress the growth defect without altering tRNA levels.

The finding that *S. pombe tan1Δ* mutants are temperature sensitive due to RTD is consistent with our work in *S. cerevisiae* showing that *tan1*Δ mutants are temperature sensitive due to RTD (Chernyakov et al. 2008; Kotelawala et al. 2008; Dewe et al. 2012)(Fig. S6), and emphasizes the conservation of this aspect of Tan1 biology across 600 million years (Parfrey et al. 2011). Recently, several individuals with a specific syndromic neurodevelopmental disorder were found to have bi-allelic mutations in the human *TAN1* ortholog *THUMPD1*, which was linked to complete loss of ac^4^C in small RNA and in purified tRNA^Ser(CGA)^ for one set of derived lymphoblastoid cell lines, and for each of two human KO cell lines (Broly et al. 2022). Based on our results, one plausible explanation for the neurological disorder of patients with bi-allelic *THUMPD1* mutations is that one or more tRNAs is destabilized due to the loss of ac^4^C, resulting in degradation of those tRNAs by the RTD pathway.

We interpret the results presented here as an expansion of our prior model to explain the common link between the RTD pathway and GAAC activation that we observe in both *S. pombe trm8Δ* mutants and *S. cerevisiae trm8Δ trm4Δ* mutants (Fig. 8). In the original model (De Zoysa and Phizicky 2020), increased temperature in both of these mutant strains concomitantly reduces the growth rate and triggers decay of specific hypomodified tRNAs (tRNA^Tyr(GUA)^ and tRNA^Pro(AGG)^ in *S. pombe trm8Δ* mutants, and tRNA^Val(AAC)^ in *S. cerevisiae trm8Δ trm4Δ* mutants), which in turn activates the GAAC pathway. In *S. pombe trm8Δ* mutants, GAAC activation results in further loss of the tRNA^Tyr(GUA)^ and tRNA^Pro(AGG)^, explaining why mutation of GAAC components suppresses their temperature sensitivity. In contrast, in *S. cerevisiae trm8Δ trm4Δ* mutants, GAAC activation prevents further loss of the tRNA^Val(AAC)^, explaining why mutation of GAAC components exacerbates their temperature sensitivity and loss of tRNA^Val(AAC)^.

**Fig. 8.**
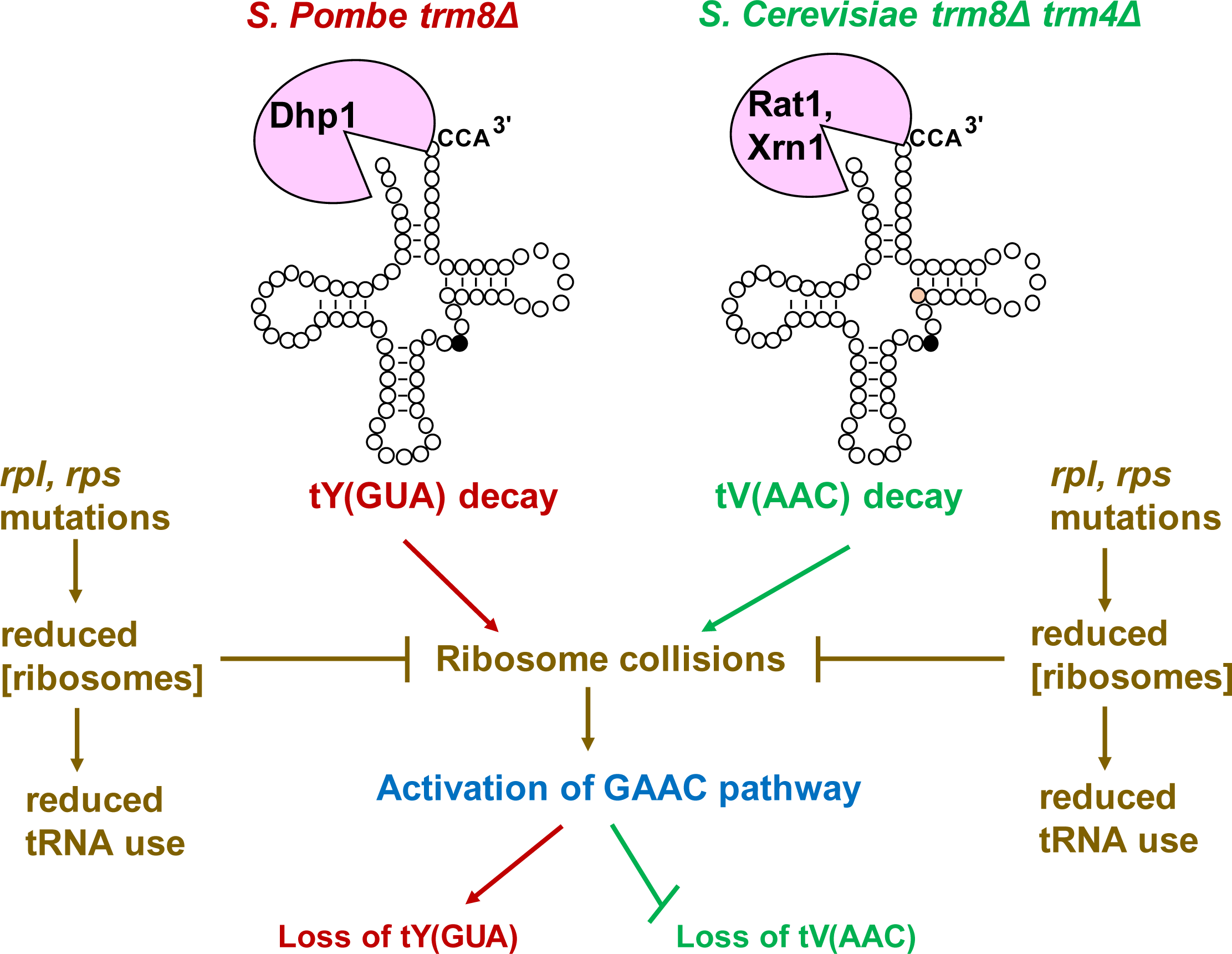
A model describing the roles of *rpl*, *rps* mutations in the biology of *S. pombe trm8Δ* and *S. cerevisiae trm8Δ trm4Δ* mutants. In both *S. pombe trm8Δ* mutants (red) and *S. cerevisiae trm8Δ trm4Δ* mutants (green), elevated temperature (38.5°C and 32°C respectively) triggers RTD of the corresponding tRNA (decay of tRNA^Tyr(GUA)^ by Dhp1/Rat1 in *S. pombe* and decay of tRNA^Val(AAC)^ by Rat1/Dhp or Xrn1 in *S. cerevisiae*). In each organisms, the reduced levels of the corresponding tRNA result in increased ribosome collisions and activation of the GAAC pathway (blue), leading to further loss of tRNA^Tyr(GUA)^ in *S. pombe* and prevention of further loss of tRNA^Val(AAC)^ in *S. cerevisiae* (De Zoysa and Phizicky 2020). In both organisms, the *rpl* and *rps* mutations (brown) result in a reduced concentration of ribosomes, leading to inhibition of ribosome collisions and reduced GAAC activation. In both organisms, the reduced concentration of ribosomes is also expected to result in reduced translation and reduced demand for tRNA use in translation.

In the expanded model (Fig. 8), the *rpl* and *rps* mutations lead to a reduced concentration of ribosomes in both *S. pombe* and *S. cerevisiae*, consistent with the fact that many *S. cerevisiae rpl* or *rps* mutants result in reduced concentrations of 60S or 40S subunits and reduced translation (Warner 1999; Cheng et al. 2019). The reduced population of ribosomes has two consequences. First, there are reduced collisions of translating ribosomes, known to result in reduced activation of the GAAC (integrated stress response) pathway in *S. cerevisiae* and mammals (Wu et al. 2020; Yan and Zaher 2021; Kim and Zaher 2022). This effect is seen in *S. pombe trm8Δ* mutants and *S. cerevisiae trm8Δ trm4Δ* mutants, leading to suppression of the temperature sensitivity of *S. pombe trm8Δ* mutants and exacerbation of the temperature sensitivity of *S. cerevisiae trm8Δ trm4Δ* mutants. Second, the reduced population of ribosomes in strains with *rplΔ* and *rpsΔ* mutations results in reduced translation and therefore a reduced requirement for tRNA. This effect is seen in *S. pombe tan1Δ* mutants, in which an *rpl502Δ* mutation does not restore tRNA^Leu(AAG)^ levels at 39°C, does not significantly inhibit the relatively weak GAAC activation observed in *tan1Δ* mutants, but suppresses the *tan1Δ* growth defect more efficiently than a *gcn2Δ* mutation.

We infer that the ribosomal protein mutations result in a reduced number of ribosomes, rather than resulting in a specific mode of ribosome function, for two reasons. First there is large diversity in the *rpl* and *rps* mutations/deletions that suppress the growth defect of *S. pombe trm8Δ* mutants (encoding seven different large subunit proteins and two different small subunit proteins) or of *S. pombe tan1Δ* mutants (encoding four different large subunit proteins and one small subunit protein, including one from each subunit that was not observed or tested in *trm8Δ* mutants). Second, these eight different proteins of the large subunit and three different proteins of the small subunit are found at different locations on the mature ribosome (Ben-Shem et al. 2011; Yusupova and Yusupov 2014). The isolation of the *grn1-N37fs* mutation as a *tan1Δ* suppressor also implicates the concentration of the 60S subunit, given its known crucial role in 60S biogenesis (Bassler et al. 2001; Du et al. 2006).

It is curious that many of the spontaneous *S. pombe trm8Δ* and *tan1Δ* suppressors that we sequenced after screening for 3-AT resistance and 5-FU sensitivity had mutations in *rpl* genes or in the *grn1* gene (implicated in 60S production), but not in *rps* genes. Although we do not know the source of this bias, results of others show distinct differences in the growth rates of individual *S. cerevisiae rplΔ* or *rpsΔ* mutant strains, and significant differences between the properties of *rplΔ* and *rpsΔ* mutant strains (Steffen et al. 2008; Steffen et al. 2012; Cheng et al. 2019). Presumably the same is true in *S. pombe rplΔ* or *rpsΔ* mutant strains. We note in this connection that different tested *S. pombe rplΔ* or *rpsΔ* mutations have different growth properties in 3-AT.

According to the proposed model, the mechanism by which *rpl* or *rps* mutations suppress will depend on which specific consequence of reduced ribosome concentration is more important in the particular modification mutant. In *S. pombe trm8Δ* mutants, the GAAC activation is large and concomitant with the onset of a growth defect and the initial loss of tRNA, and the *rpl* mutations unambiguously suppress the GAAC activation, whereas in *S. pombe tan1Δ* mutants, the GAAC activation is not as large, and suppression occurs more because of the reduced need for tRNA. As both the reduced GAAC activation and the reduced need for tRNA act in the same direction, to suppress the growth defect in *S. pombe*, the dominant effect of the *rpl* mutation (inhibition of GAAC activation or reduced need for tRNA) will depend on the modification mutant, and on the specific conditions of the growth experiment. The situation is more complicated in *S. cerevisiae trm8Δ trm4Δ* mutants, because the mutations in ribosomal protein genes are expected to exert their effects in two opposite directions. Thus, in *S. cerevisiae trm8Δ trm4Δ* mutants, *rplΔ* and *rpsΔ* mutations are expected to exacerbate the growth defect by preventing GAAC activation, resulting in increased loss of tRNA (Fig. 8), and are also expected to suppress the growth defect by reducing tRNA demand. The result of these two opposite effects is a more modest predicted effect of *rplΔ* and *rpsΔ* mutations on the growth of *S. cerevisiae trm8Δ trm4Δ* mutants than on the growth of *S. pombe trm8Δ* or *tan1Δ* mutants.

Based on our model, it seems likely that mutations in *rpl* or *rps* genes will similarly suppress the growth defect of other *S. pombe* modification mutants, particularly those in which the growth defect is due to RTD of an elongator tRNA. Moreover, as the connection between the ribosome and body modification mutants extends to *S. cerevisiae trm8Δ trm4Δ* mutants, it seems likely that similar modification mutants in other organisms, including humans will have corresponding effects.

## Materials and Methods

### Yeast strains

*S. pombe* haploid WT and two independent *S. pombe trm8Δ::kanMX* strains were derived from SP286 (*ade6-M210/ade6-M216, leu1-32/leu1-32, ura4-D18/ura4-D18 h+/h+*)(Kim and Johnson 1988), and were obtained from Jeffrey Pleiss (Cornell University). *S. pombe trm8Δ::kanMX* and *trm8Δ::kanMX rpl502Δ::hygMX* strains were constructed from haploid WT strains by PCR amplification of *trm8Δ::kanMX* DNA or *rpl502Δ::hygMX*, followed by linear transformation using lithium acetate (Bahler et al. 1998). Other deletion strains were constructed in the same way (Table S1). *S*. *cerevisiae* deletion strains (Table S2) were constructed by linear transformation with PCR amplified DNA from the appropriate knockout strain (Giaever et al. 2002). All *S. pombe* and *S. cerevisiae* strains were confirmed before use by PCR amplification, and two or three independent isolates were compared for growth properties before use.

### Plasmids

Plasmid AB553-1 was derived from pREP3X (De Zoysa and Phizicky 2020). The *S. pombe* plasmids expressing *S. pombe* P*rpl502+ rpl502^+^*(EAH 284-1) and *S. pombe* P*rpl1701+ rpl1701^+^* (EAH 282-1) were generated by inserting PCR amplified DNA genomic DNA (including ∼1000 bp upstream and downstream) into the Not I and Xho I sites of AB 553-1, removing the P*nmt1* promoter (Table S3).

### Yeast media and growth conditions

*S. pombe* strains were grown in rich (YES) media (containing 0.5% yeast extract, 3% glucose, and supplements of 225 mg/l of adenine, uracil, leucine, histidine and lysine), or Edinburgh minimal media complete (EMM-C) containing glucose and 225 mg/l of all amino acids, adenine, and uracil, as well as 100 mg/l of para-amino benzoic acid and inositol, and 1125 mg/l of leucine for Leu^-^ auxotrophs (Udagawa et al. 2008). For temperature shift experiments, cells were grown in YES or EMMC-Leu media at 30°C to OD600 ∼ 0.5, diluted to ∼ 0.1 OD in pre-warmed media and grown for 9-10 hours as indicated (De Zoysa and Phizicky 2020).

### Bulk RNA preparation and northern blot analysis

For northern analysis, 3 biological replicates were grown in parallel, and then bulk RNA was isolated from pellets derived from 2 ml culture using acid washed glass beads and phenol, separated on a 10% polyacrylamide (19:1), 7M urea, 1X TBE gel, transferred to Amersham Hybond-N^+^ membrane, and analyzed by hybridization to 5’ ^32^P-labeled DNA probes (Table S4) as described ((Alexandrov et al. 2006).

### Isolation and purification of tRNA, and analysis of nucleoside modifications

*S. pombe tan1Δ* mutants and WT cells were grown to OD ∼ 1.0 in YES media at 30°C. Then bulk RNA was extracted from ∼ 300 OD of pellets with hot phenol, and tRNA^Leu(CAA)^ was purified using the 5’-biotinylated oligonucleotide ONV22 (5’ TGGTGACCAGTGAGGGATTCGAAC, complementary to residues 76-53, using standard nomenclature), digested to nucleosides, and analyzed by HPLC as previously described (Jackman et al. 2003).

### Quantitative RT-PCR analysis

Strains were grown in triplicate to log phase and bulk RNA was prepared from 2-5 OD pellets using acid washed glass beads and phenol, treated with RQ1 RNase-free DNase (Promega), reverse transcribed with Superscript II Reverse Transcriptase, and cDNA was analyzed by quantitative PCR as described (Preston et al. 2013).

### Spontaneous suppressors and whole-genome sequencing

To select spontaneous suppressors of *S*. *pombe trm8Δ* mutants, individual colonies were inoculated into YES media at 30°C and ∼10^7^ cells of overnight cultures were plated YES media at 38°C and 39°C. Whole genome sequencing was performed by the University of Rochester Genomics Center at a read depth of 20–110 per genome nucleotide.

## Acknowledgments

We also thank Dr. Elizabeth Grayhack for valuable discussions and comments during the course of this work and for critical reading of the manuscript, Dr. Franziska Stegemann for help in manuscript preparation, and members of the Phizicky and Grayhack labs for discussions throughout this work. This research was supported by National Institutes of Health, National Institute of General Medical Sciences (NIH/NIGMS) grant GM052347 to E.M.P. and GM080669 to J.C.F.

## Supplementary Figure Legends

**Fig. S1.**
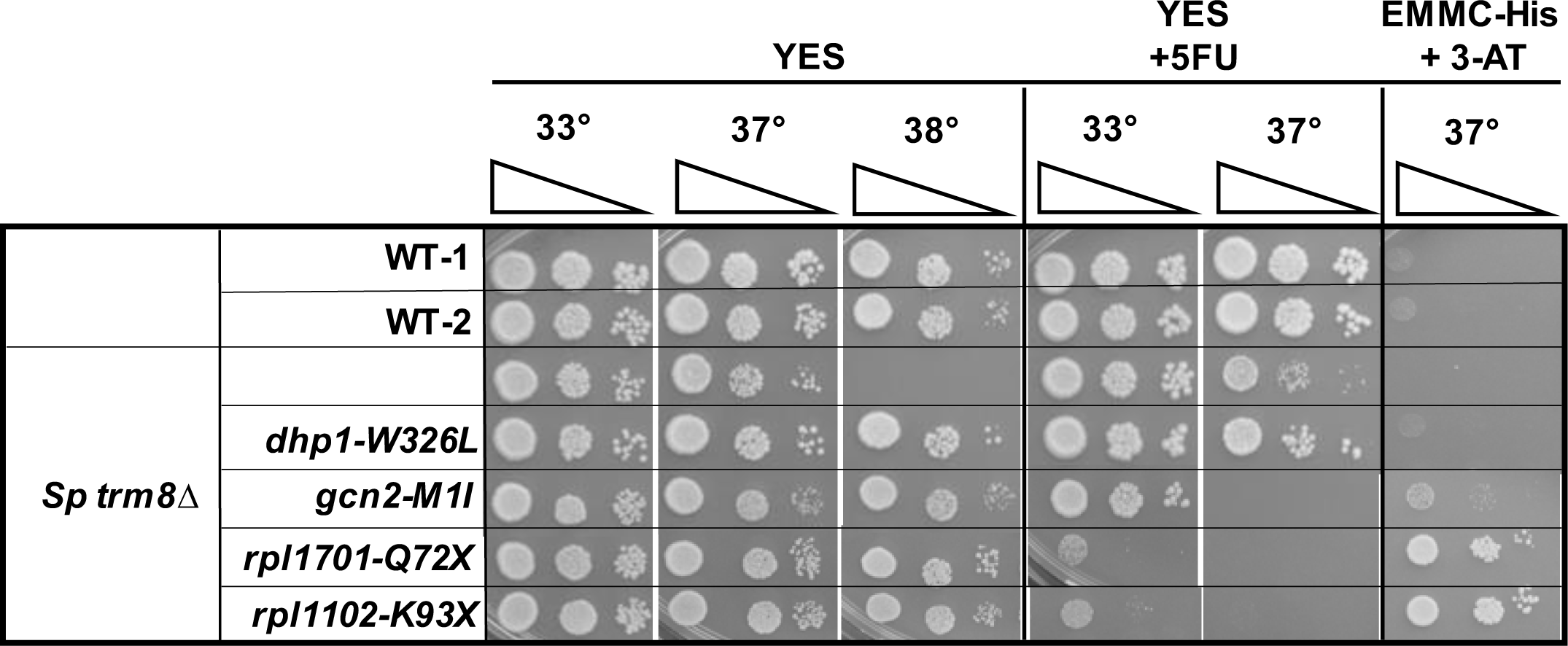
*S. pombe trm8Δ rpl1102* mutants show growth properties similar to *trm8Δ rpl 1701* mutants. *S. pombe trm8Δ* mutants and suppressors as indicated were grown overnight in YES media at 30°C and analyzed for growth on plates as described in Fig.1A.

**Fig. S2:**
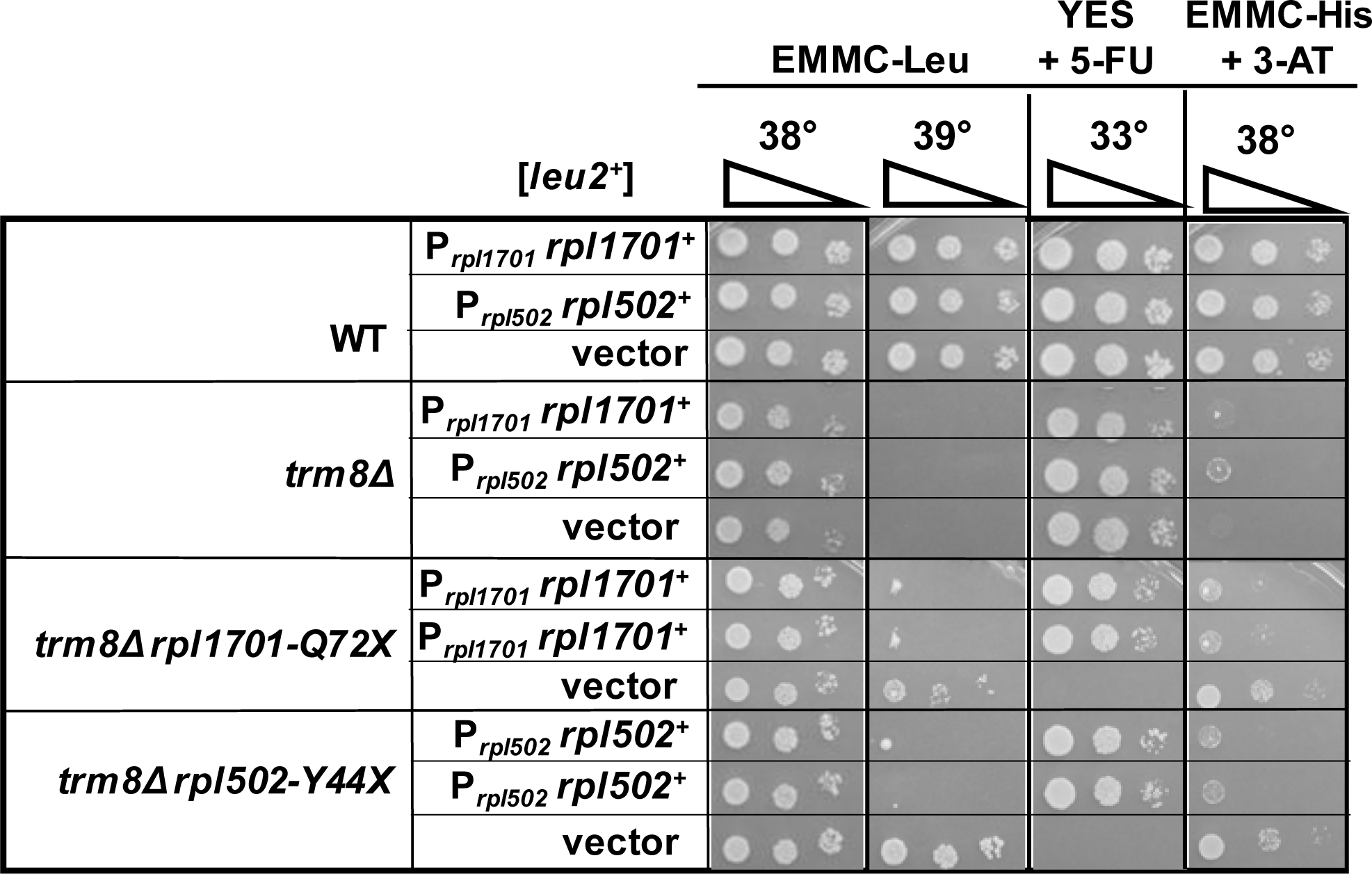
The growth properties of *trm8Δ rpl1701-Q72X* and *trm8Δ rpl502-Y44X* mutants are complemented by their respective WT genes. *S. pombe trm8Δ rpl502**-**Y44X* and *trm8Δ rpl1701-Q72X* mutants were transformed with a [*leu2^+^ rpl502*^+^] or [*leu2^+^ rpl1701*^+^] plasmid respectively, or a [*leu2^+^*] vector control, and transformants were grown overnight in EMMC-Leu media at 30 °C and analyzed for growth, as in Fig 1A, on plates containing EMMC-Leu, YES with 5-FU (30 µg/ml), and EMMC-His with 10 mM 3-AT.

**Fig. S3.**
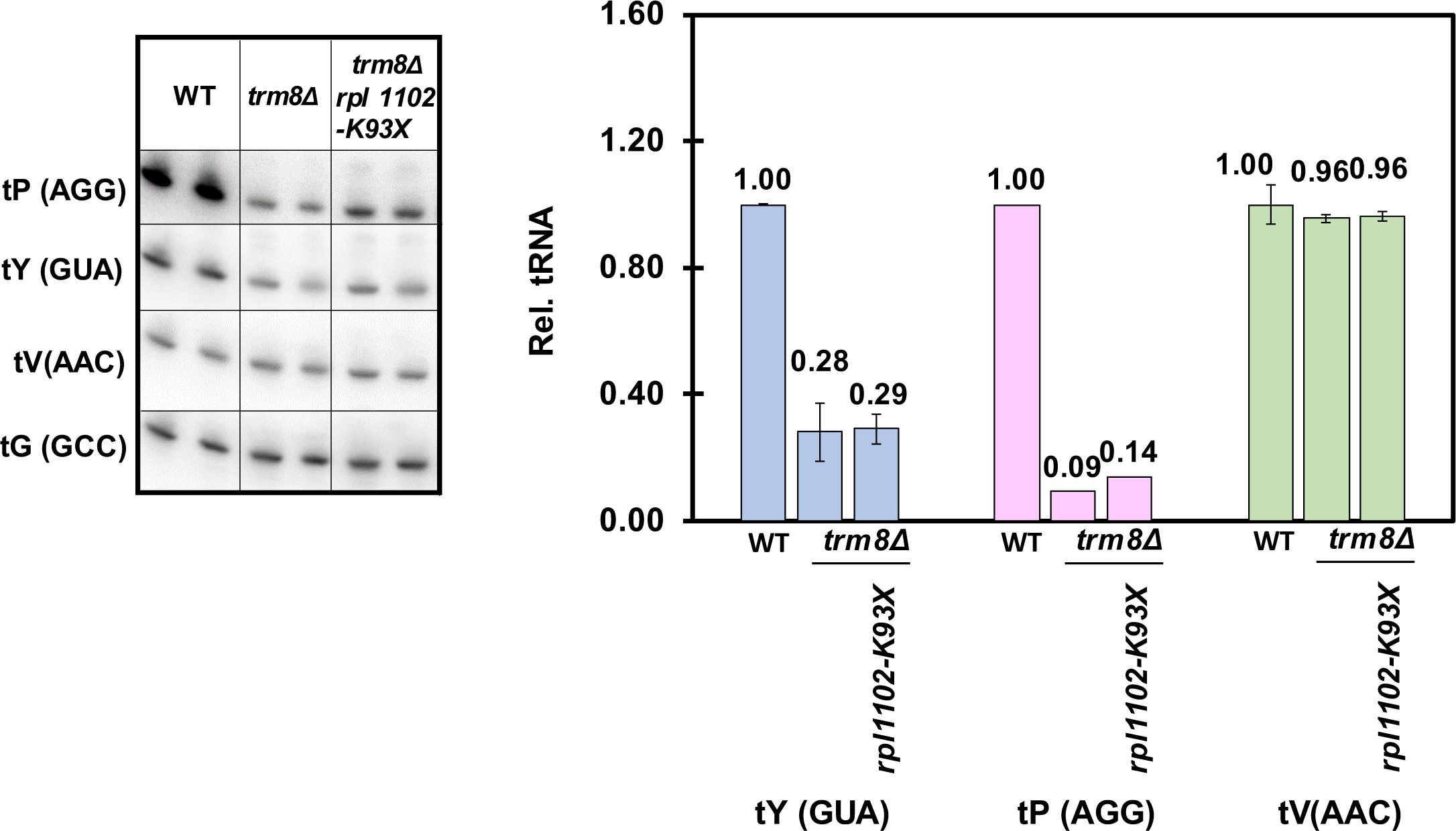
*S. pombe trm8Δ rpl1102* mutants do not significantly restore tRNA^Tyr(GUA)^ and tRNA^Pro(AGG)^ levels at 38.5 °C. Strains were transformed with a *leu2^+^*vector and then grown in EMMC-Leu media at 30°C and shifted to 38.5°C. Growth was monitored for 10 hours before harvest, and then bulk RNA was isolated and analyzed by northern blot analysis as described in Materials and Methods, and tRNA levels were quantified as described in Fig. 1C. Note that for this experiment biological replicates were compared, rather than triplicates.

**Fig. S4.**
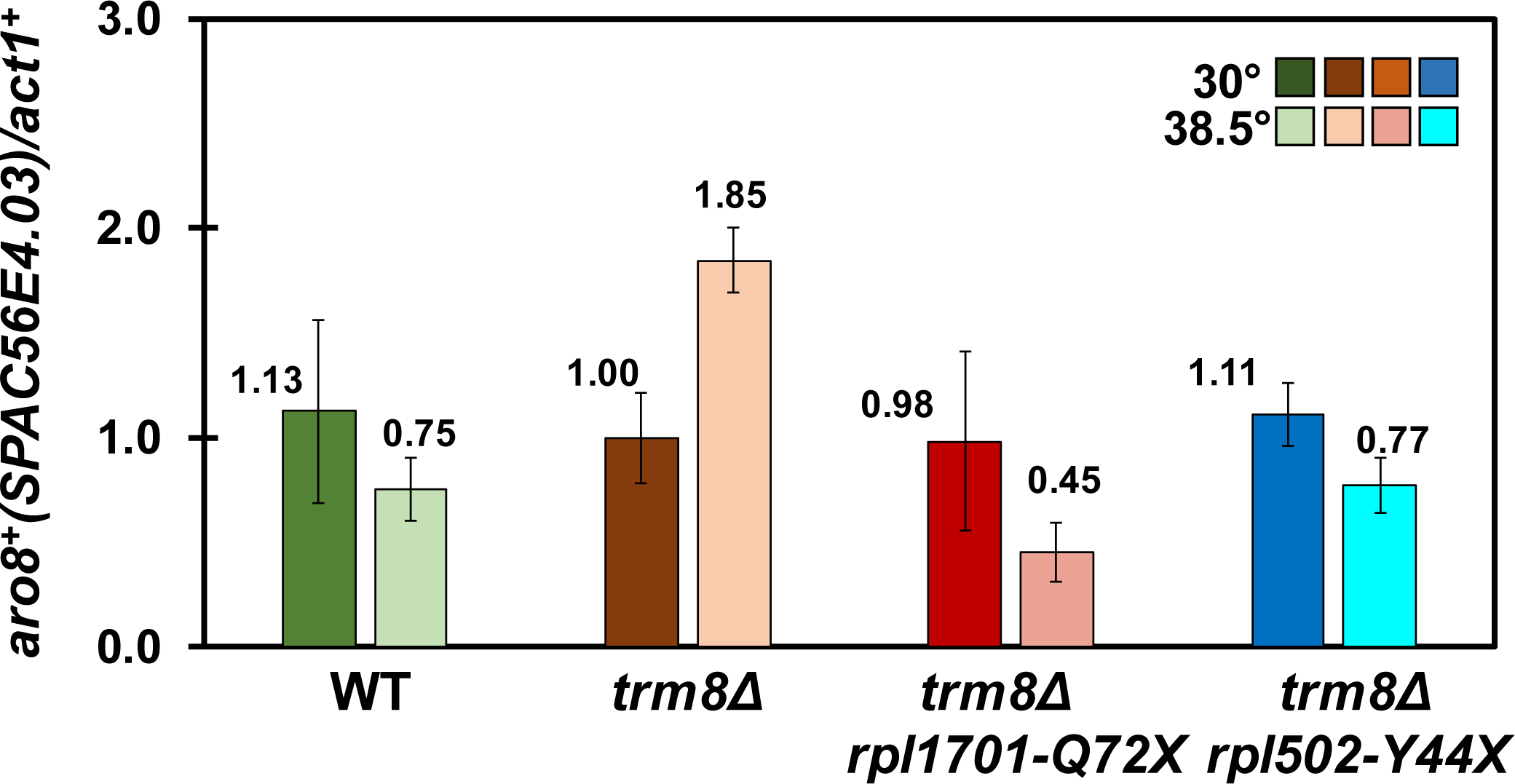
*S. pombe trm8Δ rpl1701-Q72X* and *trm8Δ rpl502-Y44X* mutants have reduced GAAC activation of *aro8^+^* mRNA at 38.5°C, relative to *trm8Δ* strains. Bulk RNA from the experiment in Fig. 1B was used to analyze *aro8^+^* mRNA levels, as described in Fig 1D.

**Fig. S5.**
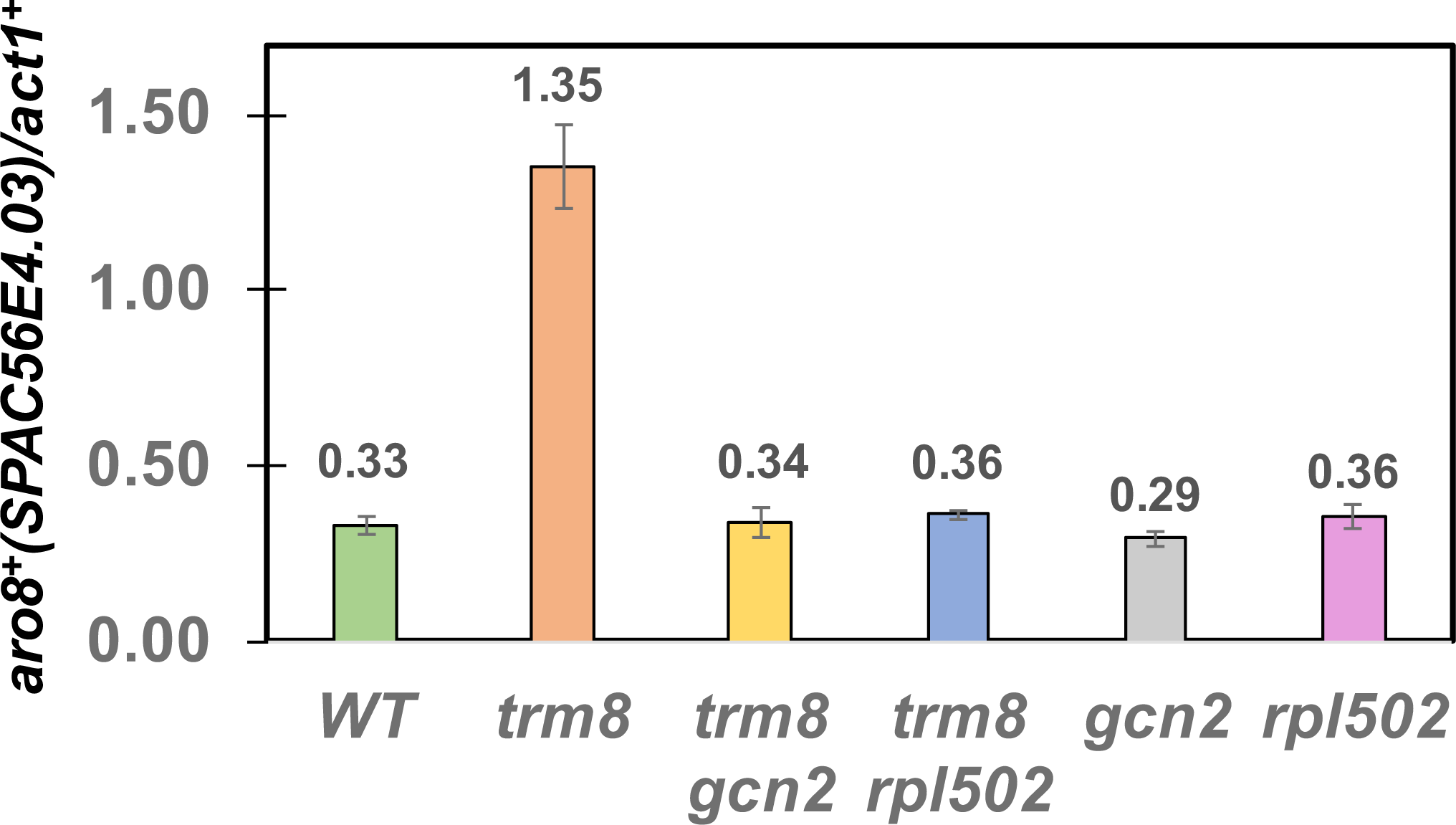
*S. pombe trm8Δ rpl502Δ* mutants have reduced GAAC activation of *aro8^+^* mRNA at 38.5°C, relative to *trm8Δ* strains. Bulk RNA from the experiment in Fig. 2B was used to analyze *aro8^+^* mRNA levels, as described in Fig 1D.

**Fig. S6.**
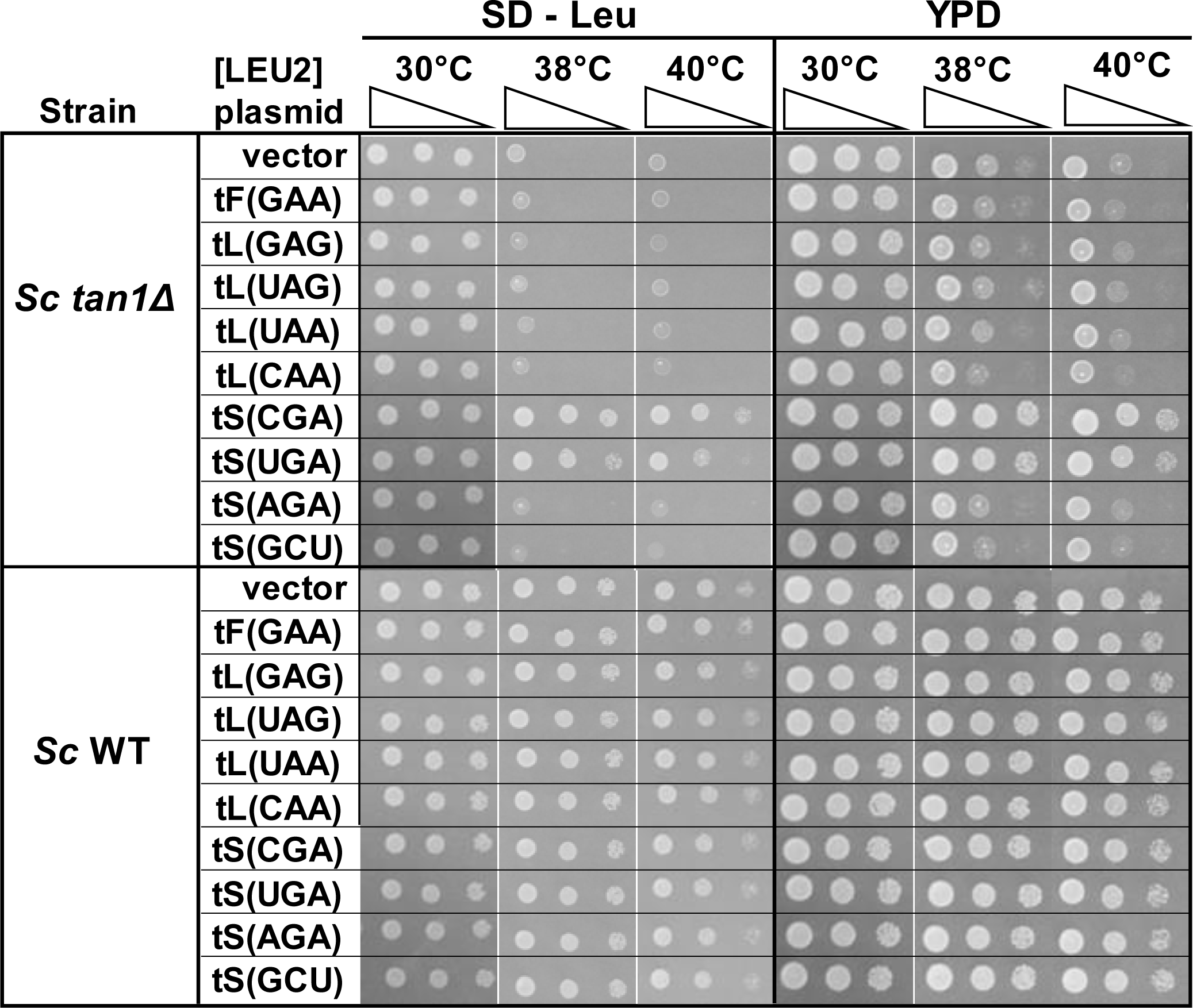
The temperature sensitivity of an *S. cerevisiae tan1Δ* mutant is efficiently suppressed by overexpression of tRNA^Ser(CGA)^ or tRNA^Ser(UGA)^. *S. cerevisiae* WT and *tan1Δ* mutants were transformed with a [*LEU2^+^*] plasmid expressing tRNAs as indicated, or a vector control, and transformants were grown overnight in SD - Leu media at 30 °C and analyzed for growth as in Fig 1A, on plates containing SD – Leu or YPD media.

**Fig. S7.**
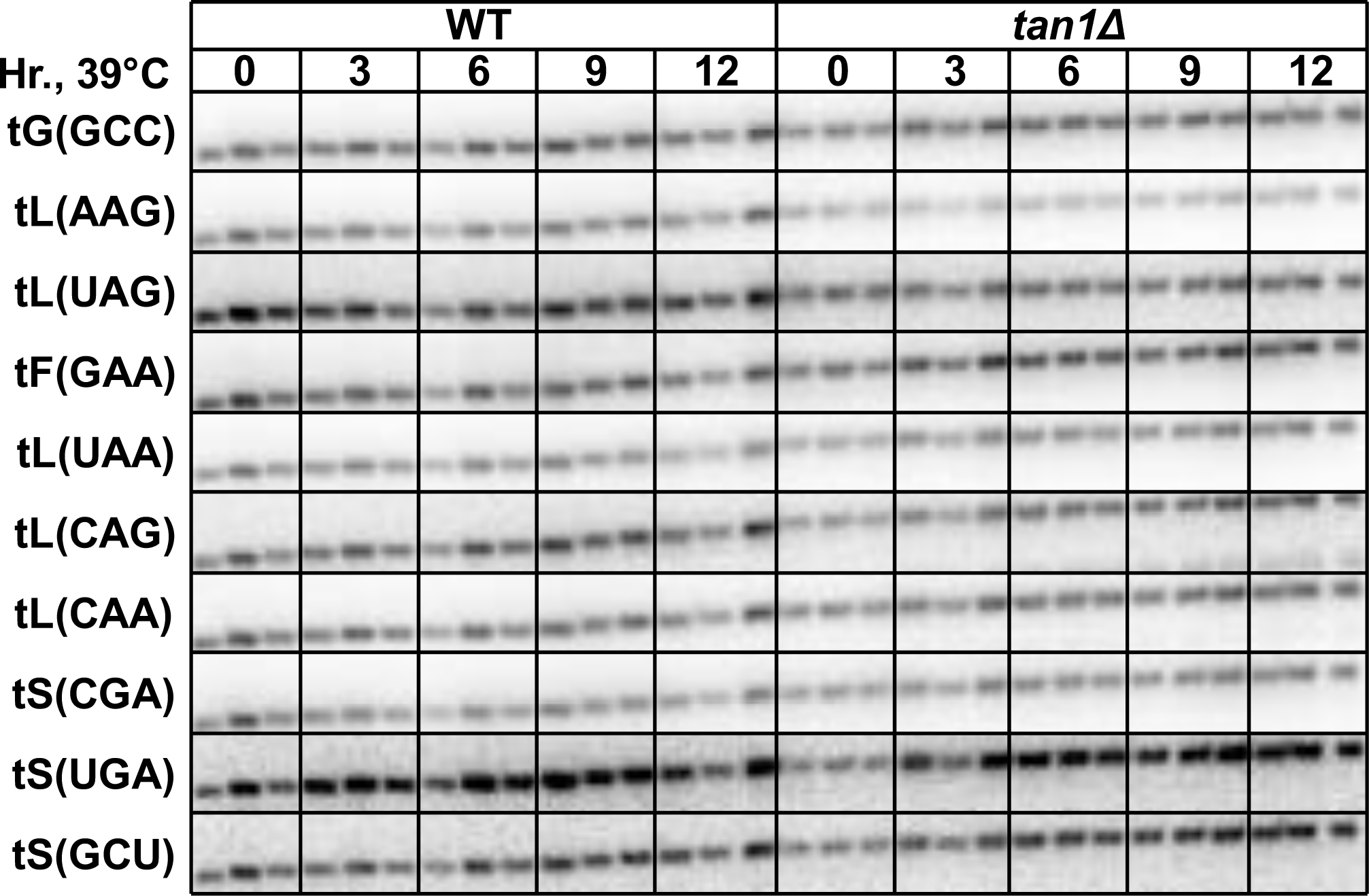
*S. pombe tan1Δ* mutants have reduced levels of tRNA^Leu(AAG)^ and tRNA^Leu(UAG)^ at 39°C. *S. pombe* WT and *tan1Δ* mutants were transformed with a [*leu2^+^*] plasmid, and transformants were grown in EMMC - Leu media to mid-log phase, diluted and shifted to 39°C and grown for 12 hours, and RNA was isolated at 0, 3, 6, 9, and 12 hours and analyzed by Northern blot as described in Materials and Methods, with the indicated probes.

**Fig. S8.**
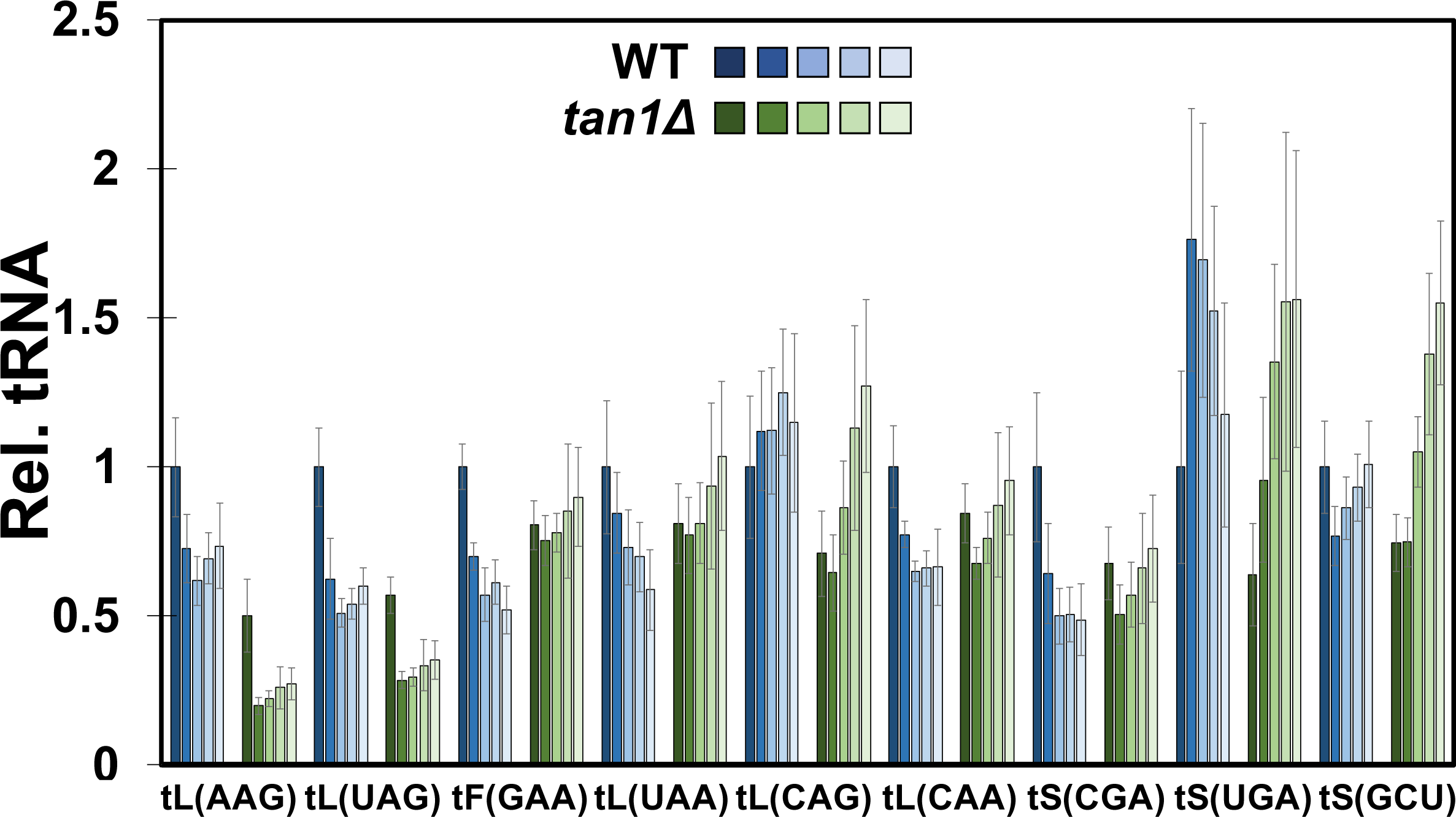
Quantification of the Northern in Fig. S7 shows that *S. pombe tan1Δ* mutants have reduced levels of tRNA^Leu(AAG)^ and tRNA^Leu(UAG)^ at 39°C. The Northern in Fig. S7 was quantified as described in Fig. 1C. Note that the data in Fig. 4C is from this quantification. shades of blue, WT strains analyzed at 0 (darkest) through 12 hours (lightest); shades of green, *tan1Δ* strains.

**Fig. S9.**
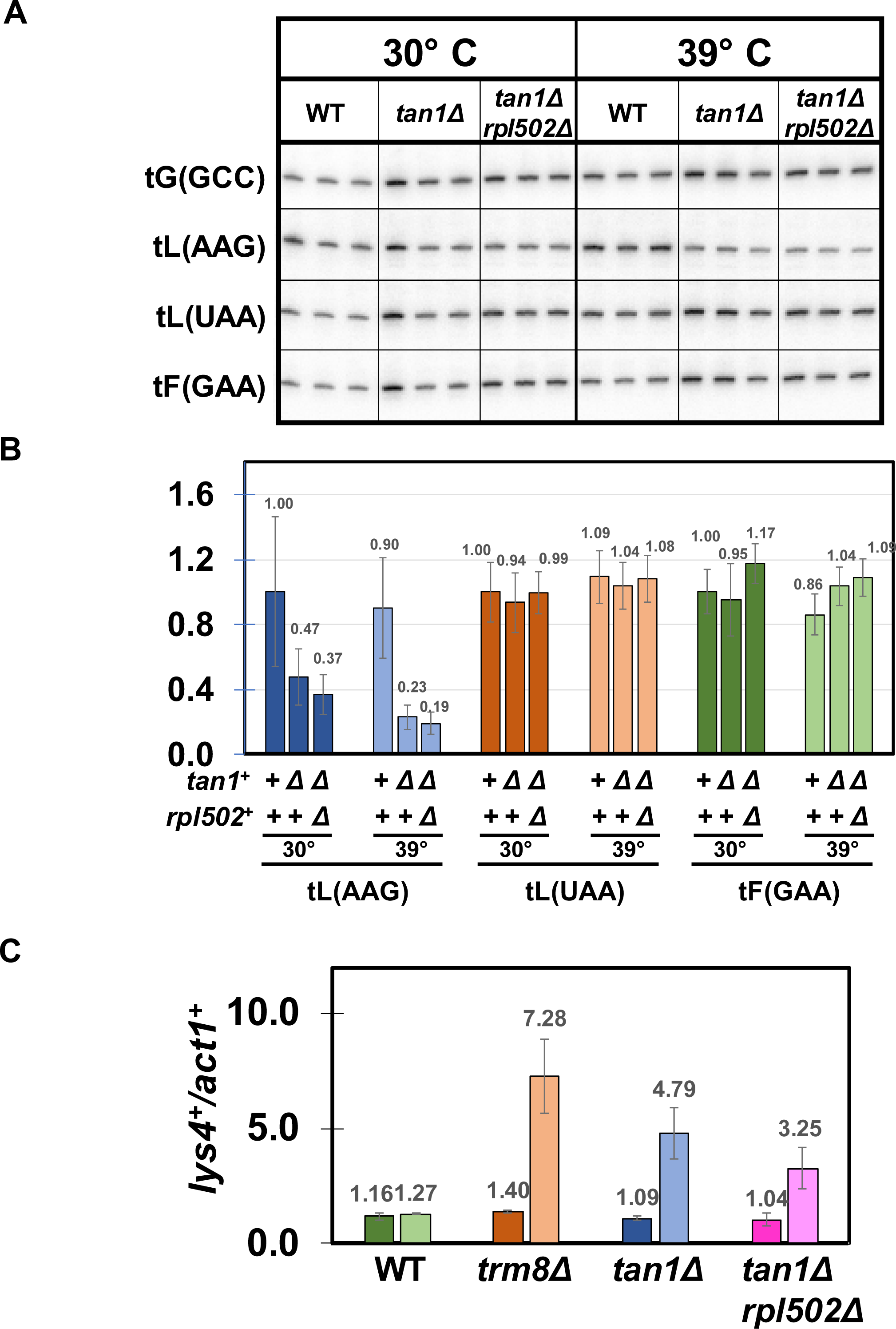
An *S. pombe tan1Δ rpl502Δ* suppressor does not rescue the tRNA^Leu(AAG)^ decay observed in *tan1Δ* mutants after growth in SD – Leu media and only minimally inhibits the modest GAAC activation. **(A). An *S. pombe tan1Δ rpl502Δ* suppressor does not rescue the tRNA^Leu(AAG)^ decay observed in *tan1Δ* mutants in SD – Leu media.** *S. pombe* WT, *tan1Δ*, and ***tan1Δ rpl502Δ*** mutants were transformed with a [*leu2^+^*] plasmid, and transformants were grown in EMMC - Leu media to mid-log phase, diluted into fresh media at 30°C and 39°C and grown for 9 hours, and then bulk RNA was analyzed for tRNA levels by northern blot analysis as described in Materials and Methods, with the indicated probes. **(B). Quantification of tRNA levels of WT, *tan1Δ*, and *tan1Δ rpl502Δ* mutants at 39°C in Fig. 6A. (C). Analysis of GAAC activation.** Bulk RNA from the growth in Fig. S9A was analyzed for GAAC activation as described in Fig. 1D.

## Supplementary Tables

**Table S1.** *S. pombe* strains used in this study.

| Strain | Parent | Genotype | Source |
| --- | --- | --- | --- |
| YTD 367-3 |  | WT | (De Zoysa and Phizicky 2020) |
| YTD 369-3 |  | *trm8Δ::kanR* (original) | (De Zoysa and Phizicky 2020) |
| YTD 556-4 | YTD 367-3 | WT *trm8Δ::kanR* (re-made) | (De Zoysa and Phizicky 2020) |
| YTD 408-1A | YTD 369-3 | *trm8Δ::kanR dhp1-W326L* | (De Zoysa and Phizicky 2020) |
| YTD 408-1E | YTD 369-3 | *trm8Δ::kanR gcn2-M1I* | (De Zoysa and Phizicky 2020) |
| YTD 375-3B | YTD 369-3 | *trm8Δ::kanR rpl502-Y44X* | this study |
| YTD 375-4H | YTD 369-3 | *trm8Δ::kanR rpl1102-K93X* | this study |
| YTD 375-4A | YTD 369-3 | *trm8Δ::kanR rpl1502-X201E* | this study |
| YTD 408-2G | YTD 369-3 | *trm8Δ::kanR rpl1701-Q72X* | this study |
| YTD 606 | YTD 556-4 | *trm8Δ::kanR rpl502Δ::HygR* | this study |
| YAH 960 | YTD 556-4 | *trm8Δ::kanR rpl1202Δ::HygR* | this study |
| YAH 954 | YTD 556-4 | *trm8Δ::kanR rpl1601Δ::HygR* | this study |
| YAH 969 | YTD 556-4 | *trm8Δ::kanR rpl1701Δ::HygR* | this study |
| YAH 961 | YTD 556-4 | *trm8Δ::kanR rpl2802Δ::HygR* | this study |
| YAH 972 | YTD 556-4 | *trm8Δ::kanR rps802Δ::HygR* | this study |
| YAH 955 | YTD 556-4 | *trm8Δ::kanR rps2801Δ::HygR* | this study |
| YTD 604 | YAH 367-3 | WT *rpl502Δ::HygR* | this study |
| YAH 957 | YAH 367-3 | WT *rpl1202Δ::HygR* | this study |
| YAH 951 | YAH 367-3 | WT *rpl1601Δ::HygR* | this study |
| YAH 964 | YAH 367-3 | WT *rpl1701Δ::HygR* | this study |
| YAH 958 | YAH 367-3 | WT *rpl2802Δ::HygR* | this study |
| YAH 967 | YAH 367-3 | WT *rps802Δ::HygR* | this study |
| YAH 952 | YAH 367-3 | WT *rps2801Δ::HygR* | this study |
| YNV 39-1 | YAH 367-3 | WT *tan1Δ::kanR* (re-made) | this study |
| YEM 71 | YNV 39-1 | *tan1Δ::kanR dhp1-I196T* | this study |
| YEM 67 | YNV 39-1 | *tan1Δ::kanR dhp1-P203L* | this study |
| YEM 46 | YNV 39-1 | *tan1Δ::kanR dhp1-P677S* | this study |
| YEM 69 | YNV 39-1 | *tan1Δ::kanR rpl1101-N7fs* | this study |
| YEM 54 | YNV 39-1 | *tan1Δ::kanR rpl1102-R133fs* | this study |
| YEM 61 | YNV 39-1 | *tan1Δ::kanR rpl1502-R67_R68_InsVR* | this study |
| YEM 43 | YNV 39-1 | *tan1Δ::kanR rpl1701-T35fs* | this study |
| YNV 57-1F | YNV 39-1 | *tan1Δ::kanR rpl2802-G113fs* | this study |
| YEM 65 | YNV 39-1 | *tan1Δ::kanR rpl3001-N80fs* | this study |
| YEM 45 | YNV 39-1 | *tan1Δ::kanR grn1-N37fs* | this study |
| YEM 55 | YNV 39-1 | *tan1Δ::kanR grn1-N37fs* | this study |
| YAH 1006 | YNV 39-1 | *tan1Δ::kanR rpl502Δ::HygR* | this study |
| YAH 1015 | YNV39-1 | *tan1Δ::kanR rpl1701Δ::HygR* | this study |
| YAH 1030 | YNV 39-1 | *tan1Δ::kanR rpl2802Δ::HygR* | this study |
| YAH 1013 | YNV39-1 | *tan1Δ::kanR rps23Δ::HygR* | this study |
| YAH 958 | YAH 367-3 | WT *rps23Δ::HygR* | this study |
| YTD 586 | YTD 367-3 | WT *gcn2Δ::HygR* | (De Zoysa and Phizicky<br>2020) |
| YAH 1005 | YNV 39-1 | *tan1Δ::kanR gcn2Δ::HygR* | this study |

**Table S2.**
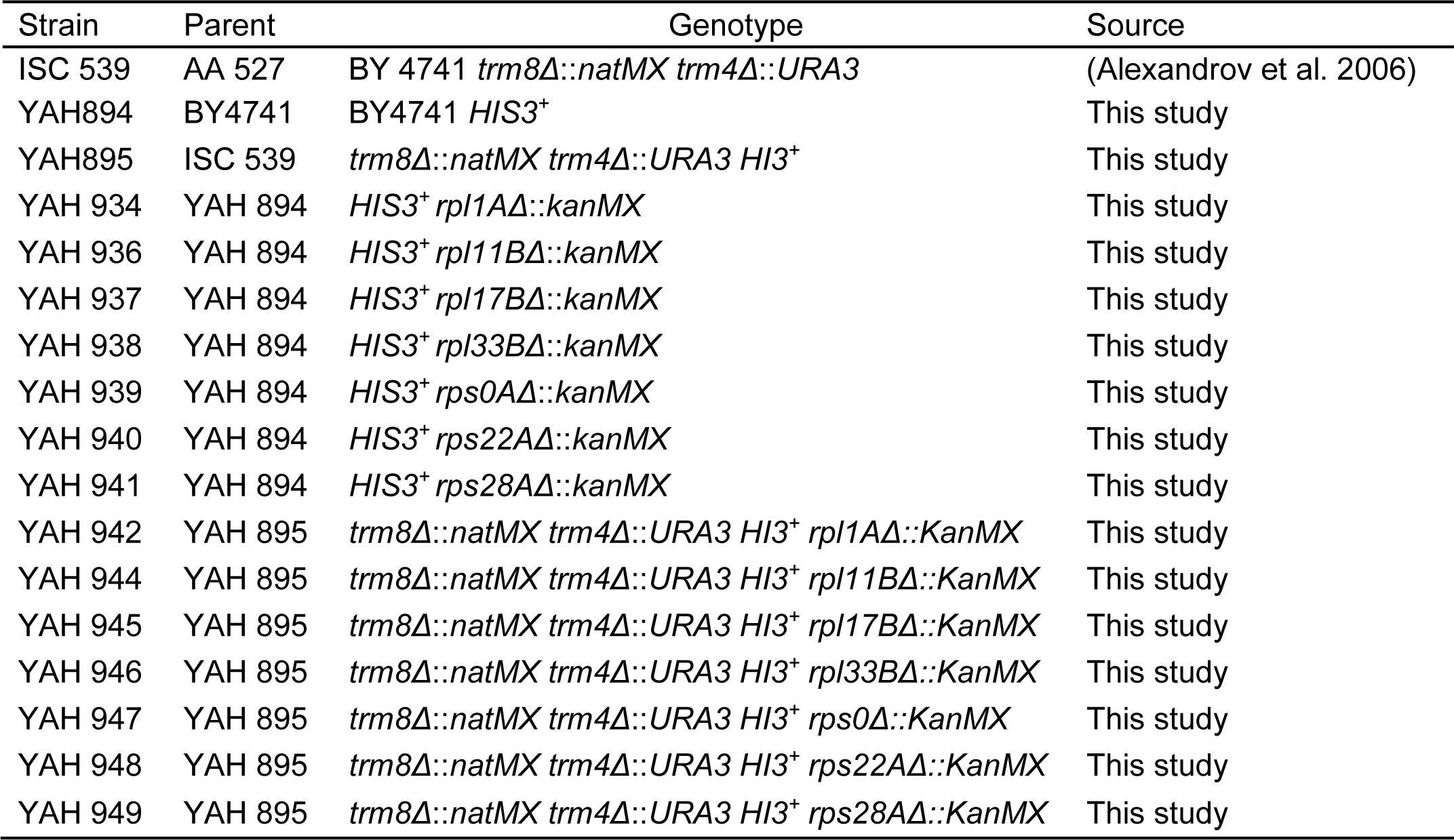
*S. cerevisiae* strains used in this study.

| Strain | Parent | Genotype | Source |
| --- | --- | --- | --- |
| ISC 539 | AA 527 | BY 4741 *trm8Δ::natMX trm4Δ::URA3* | (Alexandrov et al. 2006) |
| YAH894 | BY4741 | BY4741 *HIS3<sup>+</sup>* | This study |
| YAH895 | ISC 539 | *trm8Δ::natMX trm4Δ::URA3 HIS3<sup>+</sup>* | This study |
| YAH 934 | YAH 894 | *HIS3<sup>+</sup> rpl1AΔ::kanMX* | This study |
| YAH 936 | YAH 894 | *HIS3<sup>+</sup> rpl11BΔ::kanMX* | This study |
| YAH 937 | YAH 894 | *HIS3<sup>+</sup> rpl17BΔ::kanMX* | This study |
| YAH 938 | YAH 894 | *HIS3<sup>+</sup> rpl33BΔ::kanMX* | This study |
| YAH 939 | YAH 894 | *HIS3<sup>+</sup> rps0AΔ::kanMX* | This study |
| YAH 940 | YAH 894 | *HIS3<sup>+</sup> rps22AΔ::kanMX* | This study |
| YAH 941 | YAH 894 | *HIS3<sup>+</sup> rps28AΔ::kanMX* | This study |
| YAH 942 | YAH 895 | *trm8Δ::natMX trm4Δ::URA3 HIS3<sup>+</sup> rpl1AΔ::KanMX* | This study |
| YAH 944 | YAH 895 | *trm8Δ::natMX trm4Δ::URA3 HIS3<sup>+</sup> rpl11BΔ::KanMX* | This study |
| YAH 945 | YAH 895 | *trm8Δ::natMX trm4Δ::URA3 HIS3<sup>+</sup> rpl17BΔ::KanMX* | This study |
| YAH 946 | YAH 895 | *trm8Δ::natMX trm4Δ::URA3 HIS3<sup>+</sup> rpl33BΔ::KanMX* | This study |
| YAH 947 | YAH 895 | *trm8Δ::natMX trm4Δ::URA3 HIS3<sup>+</sup> rps0Δ::KanMX* | This study |
| YAH 948 | YAH 895 | *trm8Δ::natMX trm4Δ::URA3 HIS3<sup>+</sup> rps22AΔ::KanMX* | This study |
| YAH 949 | YAH 895 | *trm8Δ::natMX trm4Δ::URA3 HIS3<sup>+</sup> rps28AΔ::KanMX* | This study |

**Table S3.** Plasmids used in this study.

| Plasmid | Parent | Description | Source |
| --- | --- | --- | --- |
| pREP3X |  | [*LEU2* P<sub>*nmt1*</sub>] | (Forsburg 1993) |
| AB 553-1 | pREP3X | [*LEU2* with a NotI site] | (De Zoysa and Phizicky 2020) |
| EAH 282-1 | AB 553-1 | [*LEU2* P<sub>*rpl1701*</sub> *rpl1701*<sup>+</sup>] | This study |
| EAH 284-1 | AB 553-1 | [*LEU2* P<sub>*rpl502*</sub> *rpl502*<sup>+</sup>] | This study |
| ESMO 2-1 | AB 553-2 | [*LEU2* tL(*UAG*)] | This study |
| ESMO 14-2 | AB 553-1 | [*LEU2* tL(*AAG*)] | This study |
| ESMO 8 | AB 553-2 | [*LEU2* tL(*AAG*) tL(*UAG*)] | This study |

**Table S4.**
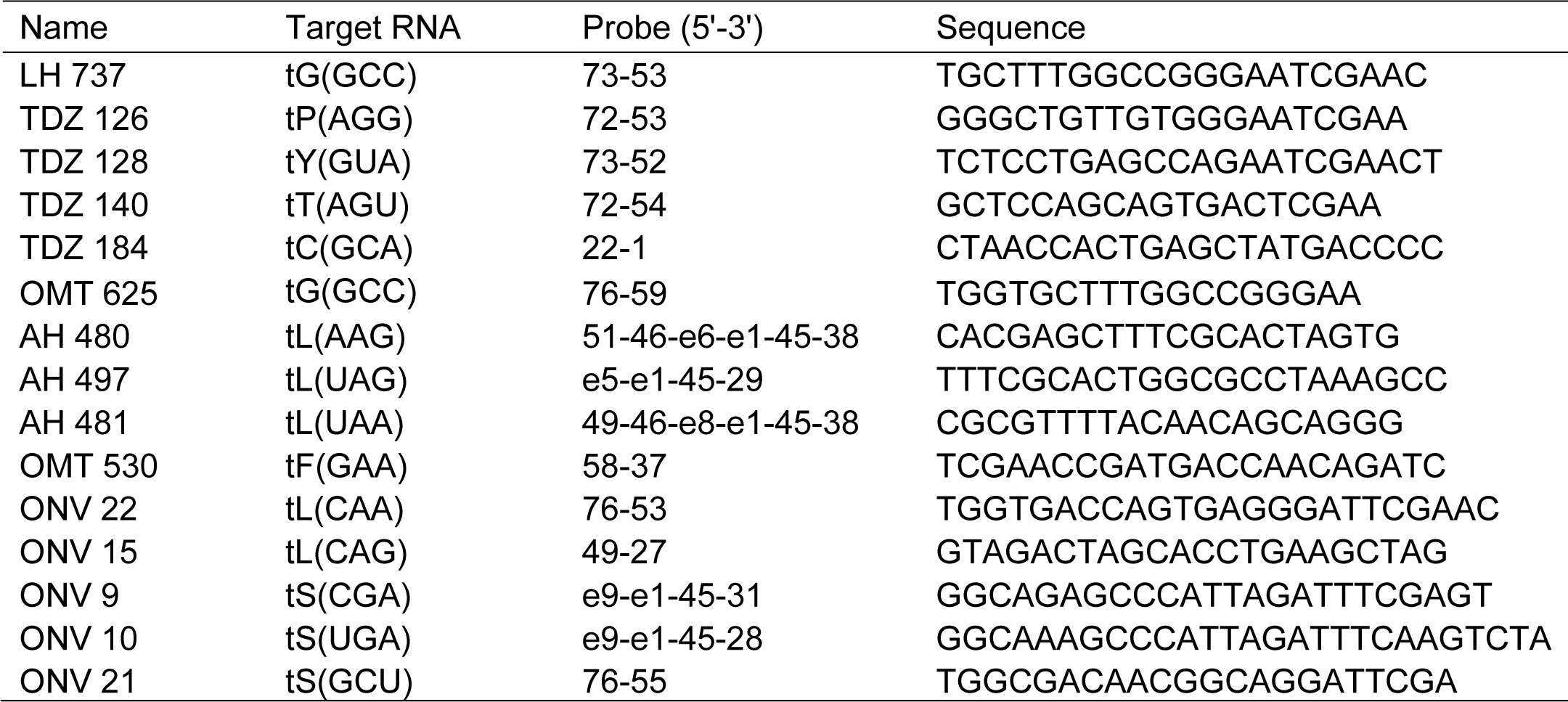
Oligonucleotides used in this study.

## Notes

### Competing Interest Statement

The authors have declared no competing interest.

